# Early Proteasome Gene Downregulation And Impaired Proteasomes Function Underlie Proteostasis Failure In Alzheimer’s Disease

**DOI:** 10.1101/2025.01.21.634128

**Authors:** Shan Jiang, Malavika Srikanth, Rossana Serpe, Shadi Yavari, Palavi Gaur, Galen Andrew Collins, Rajesh Soni, Vilas Menon, Natura Myeku

## Abstract

Alzheimer’s disease (AD) is characterized by the accumulation of pathogenic proteins, notably amyloid-beta and hyperphosphorylated tau, which disrupt neuronal function and contribute to cognitive decline. Although proteotoxic stress is well-established in AD, the role of the ubiquitin-proteasome system (UPS) in maintaining neuronal proteostasis, and how it becomes compromised during disease progression remains incompletely understood.

Here we integrated multiple approaches to characterize proteasome function, composition, and regulation in post-mortem human AD brain tissue compared to age-matched controls. These included proteasome kinetic assays, affinity purification of intact 26S proteasomes, in-gel activity assays and proteomics. According to Braak staging, we further interrogated bulk RNA-seq and single-nucleus RNA-seq (sn-RNA-seq) datasets spanning the progression of AD pathology. Finally, we examined Nrf1/NFE2L1 binding and subcellular localization to understand the transcriptional regulation of proteasome genes in AD.

We found that proteasome activity is significantly impaired in AD brains, affecting both 26S and 20S complexes. This reduction in proteolytic capacity persisted after proteasome purification, implicating intrinsic defects within the proteasome complex. Proteomic profiling revealed diminished abundances of constitutive proteasome complexes and the co-purification of proteasomes with aggregation-prone substrates (e.g., tau, α-synuclein), suggesting proteasome entrapment in pathological aggregates. Transcriptomic analyses showed progressive downregulation of constitutive proteasome subunit genes in individuals along the Braak stage axis, with downregulation apparent even at the earliest Braak stages, in tissue without overt tau aggregation. Neurons were disproportionately affected, whereas non-neuronal cells did not show substantial differences in proteasome-related gene expression, possibly through immunoproteasome induction. Despite elevated NFE2L1 expression, a key transcription factor normally driving proteasome gene transcription, AD brains exhibited impaired Nrf1 nuclear localization, preventing the expected compensatory upregulation of proteasome components.

Collectively, our findings suggest that proteasome dysfunction in AD arises early and deepens over the disease course. Intrinsic alterations in proteasome complexes, coupled with early transcriptional downregulation of proteasome subunits and disrupted Nrf1-mediated regulatory pathways, contribute to a vicious cycle of proteotoxic stress and neuronal vulnerability. Restoring proteasome function and enhancing Nrf1-driven transcriptional responses may represent promising therapeutic strategies to preserve proteostasis and mitigate neurodegeneration in AD.

## Introduction

AD is a progressive neurodegenerative disorder characterized by the pathological accumulation of two hallmark proteins: amyloid-beta (Aβ) and tau. Aβ peptides accumulate extracellularly to form amyloid plaques, while tau proteins become hyperphosphorylated and aggregate intracellularly as neurofibrillary tangles^1^. These aggregates manifest prominently in regions critical for cognition and memory, such as the hippocampus and neocortex, eventually disrupting neuronal communication, inducing synaptic loss, and leading to widespread neurodegeneration and dementia^2^. Less frequently highlighted is the fact that ubiquitin is a major component of these intracellular aggregates^3^, suggesting that dysregulated ubiquitin– proteasome-mediated degradation may be either a downstream result of disease pathogenesis or an active contributor to it.

Central to maintaining neuronal proteostasis, the delicate equilibrium of protein synthesis, folding, and degradation is the UPS. The UPS selectively degrades regulatory, short-lived and misfolded proteins through ubiquitin tagging and subsequent proteolysis by the 26S proteasome^4–6^. This ATP-dependent, ∼2.5 MDa macromolecular machinery consists of a 20S core particle (CP) harboring proteolytic sites, capped by one or two 19S regulatory particles (RP) that recognize, unfold, and translocate substrates into the CP^7^. Under physiological conditions, the proteasome not only clears damaged and misfolded proteins but also shapes the abundance of critical regulatory molecules, thus exerting tight control over neuronal function and survival.

Proteasome activity is believed to decline with age in humans and other mammals^8^, rendering post-mitotic neurons particularly vulnerable to toxic protein accumulation, especially in proteotoxic conditions such as AD. A marked reduction in proteasome function could constitute an early feature of AD, tipping the balance toward pathogenic protein aggregation and accelerating disease progression. The regulation of proteasome biogenesis is tightly controlled at the transcriptional level, ensuring that proteasome subunits are expressed in a coordinated fashion to meet the cell’s degradative demands. In mammals, the transcription factor NFE2L1 (Nrf1) maintains proteasome homeostasis by binding antioxidant response elements (AREs) in promoter regions of all proteasome subunit genes and proteasome assembly chaperones^9–11^. Nrf1 starts out as an endoplasmic reticulum (ER)-anchored protein, and its translocation to the nucleus as a transcription factor is accomplished via an elaborate activation pathway^12^. Under steady-state conditions, when proteasomes are active, Nrf1 is subjected to a swift ER-associated degradation (ERAD)^9,13^, which is firstly ubiquitinated by the ER-bound HRD1 (E3 ligase) followed by retro-translocation to the cytoplasm by p97/VCP where is immediately degraded by the proteasome (half-life∼12 min). However, when proteasome capacity is diminished, cytoplasmic Nrf1 (>120kDa) is stabilized and undergoes de-glycosylation and truncation by NGLY1 (an N-glycanase) and the DDI2 protease, respectively to generate a transcriptionally active Nrf1 (<120kDA) which is then translocated to the nucleus to bind to the ARE promoter region present in all proteasome genes to induce de novo proteasomes. This “proteasome bounce-back” response is an evolutionarily conserved mechanism that ensures rapid upregulation of proteasome genes when degradation capacity is compromised^14^. Despite this adaptive bounce-back response, in AD, this compensatory mechanism may become ineffective, contributing to the accumulation of toxic proteins.

Historically, there has been little interest in the proteasome holoenzyme’s role in determining the overall output of the UPS in AD and other degenerative diseases, as the proteasome was not thought to be the rate-limiting but merely a passive peptidase within the ubiquitin-conjugating machinery². However, recent work from our group^15–18^ and others^19–21^ has shown that the proteasome is, on the contrary, a fundamental regulator of the UPS and protein homeostasis. Therefore, changes in the proteasome levels and activity could predict the outcome of protein build-up or clearance in many neurodegenerative diseases.

In this study, we sought to clarify the functional, proteomic, and transcriptomic status of proteasomes in post-mortem human AD brains compared to age-matched, non-cognitively impaired controls. By employing a combination of proteasome assays of affinity purification, kinetic degradation, in-gel activity and proteomic assays, combined with analyses of bulk and single-nucleus transcriptomic datasets, we aimed to define how proteasome activity and composition evolve with AD severity and how these changes intersect with the broader landscape of cellular proteostasis.

The results revealed a correlation between reduced substrate degradation and reduced proteasome complexes in AD and showed that proteasomes co-purify with aggregating proteins such as tau, alpha-synuclein, and p62. Suggesting that proteasomes in AD not only exhibit reduced catalytic capacity but also become entrapped within protein aggregates.

By leveraging multiple bulk RNA-seq and sn-RNA-seq datasets spanning the progression of AD according to Braak staging, we detected a pronounced and early downregulation of genes encoding the constitutive proteasome subunits. This reduction was evident from the earliest Braak stages and became more pronounced in individuals with more advanced AD pathology. Around Braak stage IV, as the expression of constitutive proteasome genes continued to decline, we observed the emergence of immunoproteasome components, an inducible variant of the proteasome typically restricted to non-neuronal cells and upregulated under inflammatory conditions^22,23^. This shift suggests that as neurons lose their intrinsic proteasomal capacity, non-neuronal cells, such as glia, may attempt to compensate by expressing immunoproteasomes, likely in response to the heightened inflammatory milieu characteristic of AD^24^.

To elucidate the underlying cause of the reduced proteasome content in AD, we examined the Nrf1, a key transcription factor that orchestrates proteasome gene expression^9,11^. By fractionating nuclear and cytosolic compartments, we found that Nrf1 levels were diminished in the nucleus but elevated in the cytosol in AD brains. This pattern indicates that Nrf1, accumulates in the cytoplasm instead of translocating into the nucleus to initiate transcription. Impaired nuclear translocation of Nrf1 would, therefore, lead to diminished transcriptional activation of proteasome genes, contributing to the observed loss of proteasome subunits and ultimately impairing proteasomal function.

Taken together, our findings indicate that proteasome gene expression declines at the earliest stages of AD, setting the stage for diminished proteasome function. This early transcriptional deficit likely contributes to the buildup of aggregation-prone proteins, exacerbating proteotoxic stress. As these pathogenic proteins accumulate, they may further compromise proteasome activity, establishing a negative feedback loop of neuronal dysfunction and progressive neurodegeneration. Accompanied by transcriptional downregulation of Nrf1 signaling, these findings provide new insights into the molecular underpinnings of proteostasis collapse in AD and underscore the potential for therapeutic strategies aimed at bolstering proteasome activity, improving Nrf1-mediated transcription, or preventing the early downregulation of proteasome subunits

## Materials and methods

### Human Post-Mortem Brain Tissues

Human post-mortem brain tissue samples were obtained from the New York Brain Bank at Columbia University Irving Medical Center (CUIMC). Complete demographic details of the cases included in this study are provided in **Supplementary Table 1**. Specimens were collected with appropriate consent at the time of autopsy and de-identified to ensure patient confidentiality, thus exempting the use of these samples from IRB review. Approximately 300 mg tissue blocks from the Brodmann area 9 (BA9) region were dissected and immediately snap-frozen at −80°C until homogenization. We used sex-balanced tissue blocks with Control n = 40 (female = 21, male = 19) samples and AD Braak V/VI n = 60 (female = 29, male = 31). Prior to analysis, all cases underwent a comprehensive neuropathological examination by CUIMC neuropathologists to establish a neuropathological diagnosis and assign a Braak stage. To facilitate region-specific analyses, each BA9 block was subsequently separated into grey and white matter samples.

### Cell lines

We utilized HEK293-derived clonal cell lines (DS1 and DS9), originally generated by Dr. Marc Diamond’s laboratory^25^, which stably express the repeat domain of 2N4R tau bearing the P301L and V337M disease-associated mutations fused to YFP (RD-P301L/V337M-YFP). The DS1 clone does not exhibit tau aggregation or associated proteotoxicity, serving as a “non-aggregating” control. In contrast, the DS9 clone consistently forms and propagates stable tau aggregates that persist indefinitely in culture without the addition of exogenous seeds. Both DS1 and DS9 cells were maintained in DMEM supplemented with 10% FBS, 100 U/ml penicillin/streptomycin, and 1% nonessential amino acids.

### Native In-Gel Proteasome Activity Assay

To assess native proteasome activity in human cortical brain tissue, approximately 50–100 mg of frozen cortical samples were homogenized in a buffer containing 50 mM Tris-HCl (pH 7.4), 5 mM MgCl2, 5 mM ATP, 1 mM dithiothreitol (DTT), and 1 mM EDTA. To stabilize proteasome enzymatic function and inhibit phosphatases, we included 10 mM NaF, 25 mM β-glycerophosphate, a phosphatase inhibitor cocktail, and 10% glycerol. Following homogenization, the lysates were centrifuged at 20,000 × g for 25 min at 4°C to clear debris. Protein concentrations in the resulting supernatants were determined using a Thermo Scientific Pierce Bradford Plus Protein Assay (PI23238, Thermo Scientific). For native PAGE, equal amounts of protein (normalized by total protein concentration) were loaded onto a 4% non-denaturing polyacrylamide gel. The gels were run at 160 V for 180 minutes on ice to maintain native enzyme structure. Post-electrophoresis, gels were incubated for 15 min at 37°C in a homogenization buffer containing 100 μM Suc-LLVY-amc (G1101, UBP Bio), a fluorogenic substrate that measures the chymotrypsin-like proteasome activity. Active proteasome complexes (e.g., 26S and 20S) were visualized under 365 nm UV transillumination and imaged using an iPhone camera. Fluorescent band intensities corresponded to proteasome activity levels and were quantified using ImageJ or similar image analysis software.

### Affinity Purification of 26S Proteasomes

Approximately 500 mg of frozen cortical tissue was homogenized in an ATP- and MgCl2-containing buffer, supplemented with mild detergents, and centrifuged at 100,000 × g for 30 min at 4°C to obtain soluble extracts enriched in proteasome complexes. The clarified supernatant was incubated with recombinant GST-UBL (2 mg/ml) and 300 μL of glutathione– Sepharose 4B resin (17075601, Cytiva) for 2 hours at 4°C. This step leverages the ubiquitin-like (UBL) domain of Rad23B fused to GST to capture 26S proteasomes without the need for tagging proteasomes. After thorough washing to remove non-specific binding, the proteasome-GST-UBL-glutathione–Sepharose slurry was applied to a 20 mL column. To elute bound proteasomes, the column was incubated with recombinant His10-ubiquitin–interacting motif (His-UIM; 2 mg/ml), which competes for proteasome binding and releases proteasomes from the GST-UBL resin. The resulting His-UIM–proteasome complexes were then incubated with Ni2+–nitrilotriacetic acid (NTA) Agarose (30210, Qiagen). Since His-UIM binds to the Ni2+ resin, proteasomes lacking the histidine tag were recovered in the flow-through. This procedure yielded a highly enriched fraction of native 26S proteasomes. The concentration of purified 26S proteasome particles was estimated by assuming a molecular weight of approximately 2.5 MDa per particle.

### Proteasome Kinetics Assay

To evaluate the kinetics proteolytic activity of proteasomes, 15 nM of purified proteasome complexes were incubated with 50 μM Suc-LLVY-amc (G1101, UBP Bio) in a suitable assay buffer (e.g., 50 mM Tris-HCl pH 7.5, 5 mM ATP, 5 mM MgCl2). The reaction was performed in a 96-well black plate and fluorescence was recorded at 380 nm excitation and 460 nm emission every 2–5 minutes over a 120-minute period on a Spark multimode microplate reader (TECAN). The initial reaction velocity (slope) was determined from the linear portion of the fluorescence increase curve, providing a measure of proteasome catalytic efficiency under the given conditions.

### Western Blot Analysis and Subcellular Fractionation

For Western blotting, human cortical tissues (∼50 mg) or cultured cells were lysed in ice-cold RIPA buffer (1M Tris-HCl pH 7.4, 0.15 M NaCl, 0.5 M EDTA, 1% NP-40) supplemented with protease and phosphatase inhibitors. Samples were sonicated (10 seconds on ice) to ensure complete lysis and centrifuged at 3,000 × g for 10 min at 4°C. Supernatants were collected, and total protein concentration was determined via the Pierce BCA Protein Assay (23228, 23224 ThermoFisher Scientific). Protein samples were adjusted to 1 µg/µL in NuPAGE LDS Sample Buffer (NP0007, Invitrogen) containing 1 M DTT. Approximately 20 µg of total protein per sample was loaded onto 4–12% Bis-Tris gels (Life Technologies; WG1402BOx10) and resolved using MOPS running buffer (NP0001). Proteins were transferred onto 0.2 µm nitrocellulose membranes (10600001, Amersham Protran) and blocked in 5% milk in TBST for 1 hour at room temperature. Membranes were incubated with primary antibodies (diluted in SuperBlock Blocking Buffer (37515, ThermoFisher)) overnight at 4°C, followed by HRP-conjugated secondary antibodies at room temperature for 1 hour. After washes, chemiluminescent signals (Immobilon Western HRP substrate, Millipore) were detected using an Azure Biosystems 600 imaging system. Band intensities were quantified with ImageJ, normalizing signals to loading controls (e.g., actin, GAPDH, or Lamin A/C) and expressed as fold changes relative to control samples.

### Subcellular Fractionation

Subcellular fractionation was performed to separate nuclear and cytoplasmic fractions from total lysates to study the localization changes of Nrf1 in control and AD samples. Samples were processed using the NE-PER Nuclear and Cytoplasmic Extraction Reagents (78835, Thermo Fisher Scientific) according to the manufacturer’s instructions. Briefly, human cortical tissue samples (50 mg) were finely minced and washed with PBS, followed by centrifugation at 500 g for 5 minutes to obtain a cell pellet. In the case of cell lines, 4×10^6 cells were washed in PBS and centrifuged at 500 g to obtain a pellet. The pellet was homogenized using a Dounce homogenizer in CER I buffer. Cytoplasmic protein extraction was performed by vortexing the homogenate for 15 seconds, incubating on ice for 10 minutes, and adding CER II buffer. After brief vortexing and 5-minute centrifugation at 16,000 g, the supernatant (cytoplasmic extract) was collected. For nuclear protein extraction, the insoluble fraction was suspended in NER buffer, subjected to intermittent vortexing over 40 minutes on ice, and centrifuged at 16,000 g for 10 minutes. The nuclear extract supernatant was then collected. All extracts were stored at -80°C until subsequent analyses. Reagent ratios (CER I:CER II:NER) were maintained at 200:11:100 μL. Nuclear and cytosolic fractions were validated by immunoblotting for Lamin A/C (nuclear marker) and GAPDH (cytosolic marker).

### Antibodies

Antibody dilutions for Western Blots are as follows: rabbit monoclonal anti-Nrf1 (1:1,000, 8052, Cell Signaling Technologies), rabbit polyclonal anti-PSMG1/PAC1 (1:1,000, 13378, Cell Signaling Technologies), mouse polyclonal anti-Rpt5/PSMC3 (1:2500, BML-PW8770, Enzo Life Sciences), mouse monoclonal anti-Lamin A/C (1:4,000, 4777, Cell Signaling Technologies), mouse monoclonal anti-GAPDH (1:8,000, 97166, Cell Signaling Technologies), mouse monoclonal anti-β-Actin (1:8,000, 3700, Cell Signaling Technologies). Primary antibodies were diluted in SuperBlock Blocking Buffer (37515, Thermo Fisher Scientific). Secondary antibodies - anti-mouse (Peroxidase AffiniPure goat anti-mouse IgG, 115-035-146, Jackson ImmunoResearch Laboratories) and anti-rabbit (Donkey anti-rabbit IgG Cross-Adsorbed secondary antibody HRP, 31458, Invitrogen) were diluted to 1:3000 in blocking buffer.

### Quantitative Proteomics

For quantitative proteomic analysis of gel-isolated or affinity-purified 26S proteasomes from grey and white matter samples of human control and AD brains, we employed Parallel Accumulation Serial Fragmentation (PASEF) -based proteomics to improve the sensitivity and selectivity of proteomics analyses. Briefly, proteins were first denatured in 0.5% sodium deoxycholate (SDC) buffer containing 100 mM Tris-HCl (pH 8.5). The samples were heated at 60°C for 20 minutes with agitation at 1000 rpm. Reduction and alkylation of cysteine residues were carried out by adding 10 mM TCEP and 40 mM CAA and incubating at 45°C for 15 minutes. Subsequently, proteins were sonicated in a water bath and cooled to room temperature before overnight digestion with a LysC/trypsin mix (enzyme-to-protein ratio of 1:50) at 37°C and 1400 rpm. Following digestion, peptides were acidified with 1% TFA, vortexed, and desalted using SDB-RPS StageTips. The cleaned peptide samples were vacuum-dried and resuspended in 10 µl of LC buffer (3% acetonitrile, 0.1% formic acid). Peptide concentrations were determined with a NanoDrop spectrophotometer, and an input of 200 ng per sample was analyzed using a timsTOF Pro2 mass spectrometer under PASEF conditions.

### Liquid Chromatography–Tandem Mass Spectrometry (LC–MS/MS)

Peptide separations were performed on a reversed-phase C18 column (25 cm × 75 µm, 1.6 µm particle size, IonOpticks) integrated with a CaptiveSpray emitter, using a flow rate of 300 nL/min over a total gradient time of 65 minutes. Mobile phase A consisted of 0.1% formic acid in water, and mobile phase B of 0.1% formic acid in acetonitrile. The gradient profile was as follows: 2% to 25% B over 35 minutes, then 25% to 40% B over the next 10 minutes, followed by 40% to 95% B over an additional 10 minutes, and finally, a 10-minute wash step at 95% B. Mass spectrometric analyses were conducted on a timsTOF Pro2 operating PASEF mode. Ion mobility was scanned from 1/K0 = 0.6 to 1.4 V·s/cm² with a ramp time of 100 ms, maintaining a 100% duty cycle. The mass range was set from m/z 100 to 1700. Instrument parameters included a capillary voltage of 1600 V, a dry gas flow of 3 L/min, and a dry temperature of 200°C. Data acquisition was configured for 10 MS/MS frames per 1.17-second duty cycle, targeting precursor charge states of 0–5. An active exclusion window of 0.4 minutes was applied, with a target intensity set at 20,000 and an intensity threshold of 2,500. Collision-induced dissociation (CID) was performed using a collision energy of 59 eV. A polygon filter was applied in both m/z and ion mobility dimensions to selectively enrich for peptide precursors and exclude singly charged background ions, thereby improving peptide identification and quantification.

### LC-MS/MS Data Analyses

Raw PASEF data files were processed in MaxQuant (v.2.4.10.0) using the Andromeda search engine with default parameters. Initial and main search mass tolerances were set to 20 ppm and 4.5 ppm, respectively. Spectra were searched against the human reference proteome (UniProt UP000005640) using trypsin specificity and allowing up to two missed cleavages. Carbamidomethylation of cysteine was designated as a fixed modification, and acetylation (protein N-terminus), oxidation (methionine), phosphorylation (serine, threonine, tyrosine), and GlyGly (lysine) were included as variable modifications. A false discovery rate (FDR) of 1% was applied at both the peptide and protein levels, and at least one unique peptide was required for protein identification. Label-free quantification (LFQ) was performed with a minimum ratio count of one to ensure robust quantitative comparisons. The LFQ intensity values and identifications generated by MaxQuant were exported for subsequent statistical analyses. Differential protein abundance was assessed using t-tests with an FDR threshold of <0.05 to identify proteins exhibiting statistically significant changes between experimental groups.

### Proteomics Data Processing and Analysis

Raw peptide intensity data from MS analyses were processed in R using the package DEP. Proteins detected with low confidence or containing excessive missing values were excluded. The remaining intensity data underwent variance-stabilizing normalization (VSN), and any remaining missing values were imputed based on a random distribution designed to approximate low-abundance signals. A linear model combined with empirical Bayes statistics (via the limma-based ‘test_diff’ function in DEP) was applied to identify proteins differentially abundant between AD and control samples. Multiple comparisons were controlled using the false discovery rate (FDR). Differential abundance analysis results of all the detected proteins were visualized with a volcano plot with logarithmic transformed P-values (vertical axis) against AD-to-Control abundance ratios (horizontal axis) for individual proteins. To visualize the UPS-related proteins at the sample level, normalized abundance data were further transformed with Z-score normalization across the samples for each protein before applying a heatmap with proteins as rows and samples as columns with the R package pheatmap.

Finally, to examine potential physical interactions between proteasome proteins and their interactors, we retrieved experimentally validated protein–protein interaction (PPI) data from the STRING database^26^. A PPI network graph was constructed in R with igraph, where nodes represent proteins and edges represent physical interactions. Nodes are colored according to their abundance changes in AD relative to controls, with blue indicating decreased abundance in AD and red/orange indicating increased abundance.

### Bulk RNA-Sequencing Data Analysis

Publicly available bulk RNA-seq datasets from the AD Knowledge Portal (MSBB, ROSMAP, and Mayo studies) were analyzed to characterize proteasome-related gene expression changes across AD progression. Raw read counts were normalized using variance-stabilizing transformations (VSN) implemented in DESeq2. Braak staging was used as a neuropathological correlate of tau-related AD progression. Multivariate ordinal regression models, controlling for variables such as RIN, sex, age, PMI, ethnicity, education, and library size (as applicable per cohort), were used to relate gene expression to Braak scores. Multiple comparisons were corrected using the false discovery rate (FDR).

To visualize changes in proteasome gene expression across Braak stages, normalized expression data were further Z-score normalized per gene. A heatmap (rows as genes, columns as Braak stages) was then generated using the R package pheatmap, where each heatmap cell (or box) represents the averaged gene expression of all samples within a particular Braak stage. For higher-level visualization of proteasome complex expression, subunit gene expression data were collapsed by averaging normalized expression values of the constituent subunits. The resulting complex-level expression data were also Z-score normalized, enabling a grouped bar plot that highlights expression trends across Braak stages.

Finally, to assess aggregated complex expression differences between AD and control samples across the three cohorts, we performed a meta-analysis of the standardized mean differences (calculated as [AD mean – Control mean]) using the R package meta. In the resulting forest plot, each horizontal line (with a center box) reflects the confidence interval (CI) and point estimate for a single study. The size of the box corresponds to the study’s weight (sample size), while the diamond indicates the overall pooled effect estimate and its 95% CI.

### snRNA-seq data Analysis

We analyzed two large snRNA-seq datasets derived from dorsolateral prefrontal cortex (DLPFC) samples ^27,28^, both accessible through the AD Knowledge Portal. To reduce false positives and effectively handle lowly expressed proteasome subunit genes, we adopted a pseudo-bulking strategy: nuclei from each cell type within each sample were aggregated, generating a gene-by-sample count matrix per cell type. These pseudo-bulk data were normalized using variance-stabilizing normalization (VSN) implemented in DESeq2. For each cell type, multivariate ordinal regression models were fitted to examine the relationship between proteasome subunit gene expression and the Braak stage. Model covariates included RIN, sex, ethnicity, education, age, postmortem interval (PMI), and library size. Multiple testing corrections were applied using the false discovery rate (FDR) to ensure robust statistical inference. To visualize the expression changes of proteasome complexes with Braak stages per cell type, subunit gene expression per cell type was collapsed to complex expression by averaging their normalized expression levels. The complex expression data per cell type were further transformed with Z-score normalization across the samples for each complex before applying a circular heatmap comprised of regular complex-expression-against-Braak heatmaps for individual cell types.

### NFE2L1 ChIP-seq data analysis

To evaluate the binding of NFE2L1 on the transcriptional regulatory regions of the proteasome genes, we analyzed a NFE2L1 ChIP-seq dataset on HepG2 cell line from the ENCODE project (https://www.encodeproject.org/experiments/ENCSR543SBE/). The NFE2L1 binding peaks were annotated to gene transcriptional regulatory regions with the R package ChIPpeakAnno. To compare the binding of NFE2L1 on the transcriptional regulatory regions of the proteasome genes with the binding on the background genes, due to the imbalanced gene counts between the two arms, we performed random sampling with replacement to get 10,000 peak samples for each arm. The distributions of the sampled binding peak signals were visualized with histograms, and the peak signal intensity difference was analyzed using an unpaired two-tailed Student’s t-test with equal variance between groups.

### NFE2L1 Resist score

To link NFE2L1 expression with proteasome expression quantitatively, we applied a linear regression model relating NFE2L1 transcript levels to the derived proteasome 26S expression as mentioned above. Before applying the model, min-max normalization was used to adjust the expression of NFE2L1 and proteasome 26S expression within the same range (from 0 to 1). Then, the absolute value of the expression difference between NFE2L1 and proteasome 26S (|E_NFE2L1 – E_P26S|), which was denoted as delta (**Δ**), was used to identify presumably normal samples. In this study, samples with **Δ** ≦ 2 were retained for model fit. After applying the linear regression model on the identified normal samples (**Δ** ≦ 2), we derived a linear equation to predict the expression of proteasome 26S with the expression of NFE2L1. Then, the NFE2L1 resist score (NRS) was computed as the difference between the predicted and the observed proteasome expression levels (E_P26S_predicted – E_P26S_observed). For the samples whose P26S levels are in line with NFE2L1-driven predictions, their NRSs remain near zero, whereas NRSs are positive for those with lower-than-expected P26S expression and negative for those with higher-than-expected P26S expression.

### Statistical analyses

Statistical analyses for figure 1, and 7 were carried out on GraphPad Prism 10 (GraphPad Software, San Diego, CA). Data were tested for normality using the D’Agostino-Pearson test or the Shapiro-Wilk test, depending on the sample size. Homogeneity of variance was determined using Levene’s test. In Fig. 1, the proteasome kinetics assays of total lysates (Fig. 1B, D) and the native in-gel proteasome activity assay for total lysates from human post-mortem gray matter samples (Fig. 1J) were analyzed by the Mann-Whitney test. The proteasome kinetics assays of purified proteasomes (Fig. 1F, H) and the native in-gel proteasome activity assay for total lysates from human post-mortem white matter samples (Fig. 1L) were analyzed using two-tailed unpaired Student’s t-tests. In Fig. 7, the immunoblotting of total lysates from human post-mortem brain tissue for Nrf1 (Fig. 7B) was analyzed by Mann Whitney test, while two-tailed unpaired Student’s t-tests were used for PSMG1 and Rpt5 (Fig. 7C, D). For immunoblotting of subcellular fractions from cell lines (Fig. 7I-K), two-way ANOVAs with Bonferroni post hoc correction were used. P values in the figures are as follows: *P < 0.05, **P < 0.01, ***P < 0.001, ****P < 0.0001, and ns, for P > 0.05.

**Figure 1.**
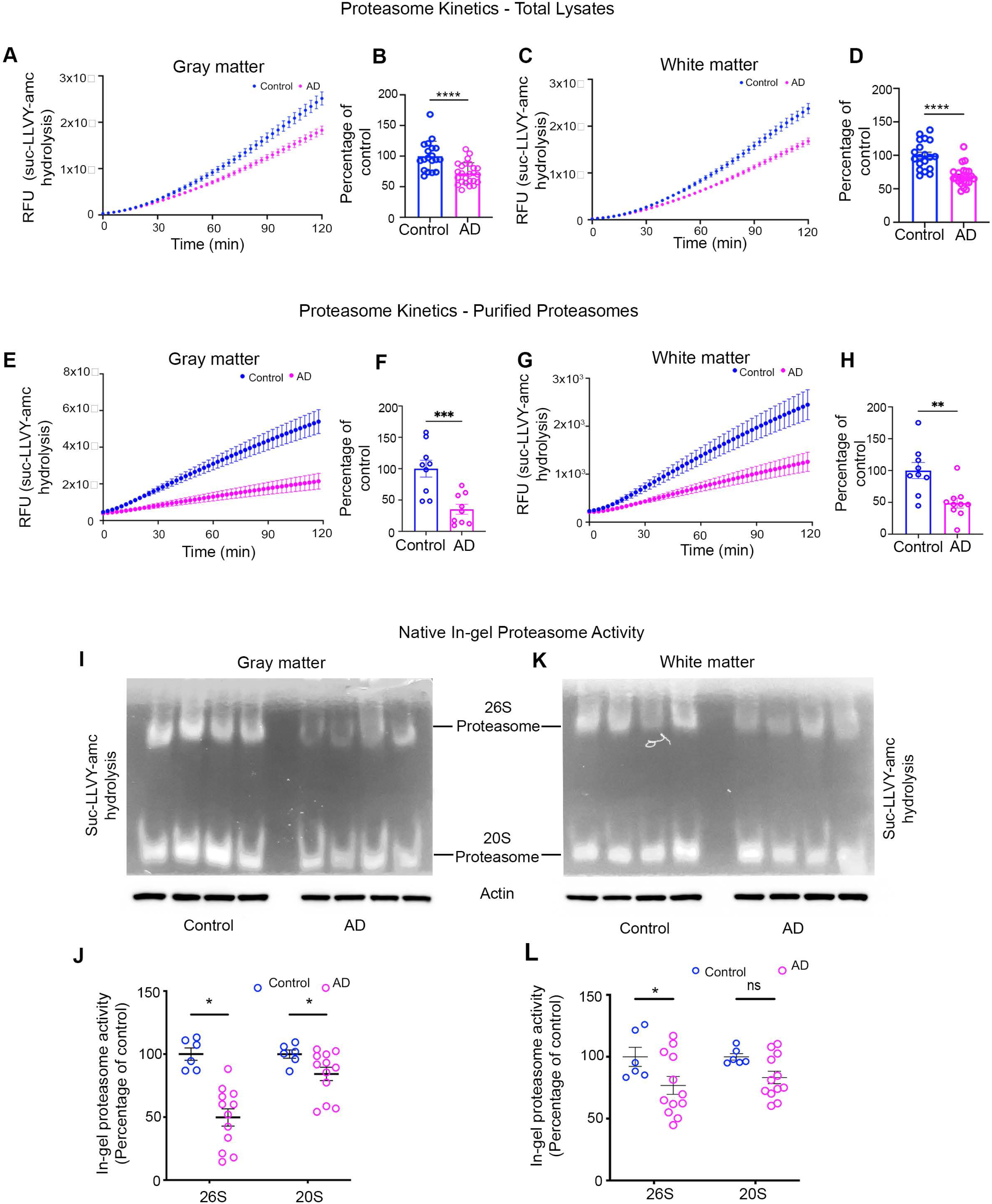
Reduced proteolytic capacity of proteasomes in AD brains. (**A–D**) Kinetic analyses of proteasome activity in total lysates isolated from (**A,B**) grey and (**C,D**) white matter. Proteasome activity was measured using the fluorogenic substrate Suc-LLVY–amc, and the rate of substrate hydrolysis was monitored over 120 minutes. (**A,C**) Representative kinetic curves of substrate hydrolysis in control (blue) and AD (pink) samples. (**B,D**) Quantification of the average proteasome activity as a percentage of control levels after two hours. Proteasome-mediated proteolysis is significantly decreased in both grey and white matter in AD. (Control n = 20, AD n = 26. Control: blue circles; AD: pink circles). (**E–H**) Kinetic analyses of purified 26S proteasomes from (**E,F**) grey and (**G,H**) white matter (control n = 9, AD n = 9). Similar to total lysates, purified proteasomes from AD brains show a marked reduction in substrate hydrolysis rate. (**E,G**) Representative kinetic curves. (**F,H**) Quantification of proteasome activity expressed as a percentage of control confirms a significant intrinsic impairment of proteolytic capacity in AD-derived proteasomes. (**I–L**) Native in-gel proteasome activity assays. (**I,K**) Representative non-denaturing gels incubated with Suc-LLVY–amc to visualize 26S and 20S proteasome activity in (**I**) grey and (**K**) white matter (control n = 6, AD n = 12). Lower panels show corresponding actin immunoblots as loading controls. (**J,L**) Quantification of proteasome activity bands expressed as a percentage of control. In (**J**) grey matter, both 26S and 20S activities are significantly reduced in AD. In **(L**) white matter only 26S activity is significantly diminished in AD, while 20S activity is not significantly affected. Each point represents an individual non-overlapping sample (control n = 35, AD n = 47) (control: blue circles; AD: pink circles). All data are presented as mean ± SEM. ns = non-significant, *p<0.05, **p<0.01, ***p<0.001, ****p<0.0001; unpaired t-tests.

### Statistical analyses for bioinformatics data

All bioinformatic analyses were performed in R. A two-tailed p-value < 0.05 was considered statistically significant unless otherwise noted. Multiple comparisons were controlled using the false discovery rate (FDR) method.

### Proteomics Analyses

Raw proteomics data were processed using the R package *DEP*. To enhance data quality, proteins not identified in all replicates of at least one condition (control or AD) were removed. A bar plot was then used to display the number of identified proteins per sample (**Supplementary Figures 1A, 2A, 3A and 4A**). Next, we performed background correction and variance-stabilizing normalization (VSN). To compare the effect of VSN, log-transformed protein abundance intensities were visualized in sample-wise box plots both before and after normalization (**Supplementary Figures 1B, 2B, 3B and 4B**).

Despite these steps, some missing data points persisted. We addressed them by imputing values from a left-shifted distribution (“MinProb”) via the *impute* function in *DEP*. The impact of imputation was assessed by plotting smoothed density curves of log-transformed protein intensities before and after imputation (**Supplementary Figures 1C, 2C, 3C and 4C**). Finally, the top 50 most variably expressed proteins were used to perform principal component analysis (PCA), the results of which were visualized in a scatter plot to inspect data structure and potential batch effects (**Supplementary Figures 1D, 2D, 3D and 4D**).

Differential abundance analysis on the processed proteomics data was performed with a protein-wise linear model combining empirical Bayes statistics using the *DEP* function *test_diff*, which is backboned by limma. The false discovery rate (FDR) approach was applied to multiple testing corrections. *P* < 0.05 was considered significant.

### Bulk RNA-Seq Analyses

To evaluate proteasome gene or complex-level expression changes across Braak stages in three large AD cohorts—Mount Sinai Brain Bank (MSBB), Religious Orders Study and Memory and Aging Project (ROSMAP), and Mayo Clinic we employed multivariate ordinal regression models (**Fig. 4A–F**):

For **MSBB** the regression model, *Braak score ∼ gene/complex expression + RNA integrity number (RIN) + sex + ethnicity + age + post-mortem interval (PMI) + library size*, was used. For **ROSMAP** the regression model, *Braak score ∼ gene/complex expression + RIN + sex + ethnicity + education + age + PMI + library size*, was used.

For the **Mayo clinic** the regression model, *Braak score ∼ gene/complex expression + RIN + sex + age + library size*, was used. Since all Mayo samples were from a white population, ethnicity was not included in this model. In these models, library size is an intermediate variable introduced by DESeq2 for normalization. FDR correction was used for multiple testing, and significance was assessed at p < 0.05.

To combine results on proteasome complex expression across MSBB, ROSMAP, and Mayo, we performed a meta-analysis of the standardized mean difference (AD_mean_ - Control_mean_) using the R package meta (**Fig. 4G**). Each study-specific effect size (with its 95% confidence interval) was pooled under a fixed-effect model, and significance was determined at p < 0.05. **snRNA-seq Analyses**

To address cell-type–specific changes in proteasome gene expression—and avoid false positives arising from cell-level analyses that treat each nucleus as an independent replicate we adopted a pseudo-bulking strategy. For each sample and each cell type c, we aggregated all nucleus-level read counts for each gene g:

For a given cell type in a given sample, pseudo-bulking for each gene was performed with the formula 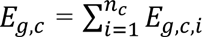, where *E_g,c_* is the single nucleus sequencing read count for gene *g* across all the nuclei of a certain cell type *c*, *n_c_* is the total cell count in the cell type *c*, and *E_g,c,i_* is the read count for gene *g* in the *i*th cell of the cell type *c*.

After pseudo-bulking for all the genes in individual samples for a certain cell type, we obtained a gene-by-sample read count matrix ***M**g×s* for the cell type.

The resulting pseudo-bulk data were variance-stabilizing normalized (VSN) in *DESeq2*, and we fit multivariate ordinal regression models similar to those above to relate proteasome subunit gene expression to the Braak stage for each cell type. The multivariate ordinal regression model was: *Braak score ∼ gene expression + RIN + sex + ethnicity + education + age + PMI + library size*, was used. The false discovery rate (FDR) approach was applied to multiple testing corrections. *P* < 0.05 was considered significant.

### NFE2L1 ChIP-Seq Enrichment and Statistical Comparison

To compare NFE2L1 (Nrf1) binding at proteasome gene promoters versus randomly selected background genes, we performed stratified random sampling with replacement to obtain 10,000 peak-signal samples per group from the NFE2L1 ChIP-seq data. Distributions of signal intensities in the two groups were evaluated using an unpaired, two-tailed Student’s t-test (assuming equal variance). Normality was assessed by the Kolmogorov–Smirnov test, and significance was set at p < 0.05.

### NFE2L1 Resist Score (NRS)

To quantify the extent of proteasome recovery through *NFE2L1* by leveraging transcriptomics, we computed the *NFE2L1* resist score (NRS) as the difference between the predicted proteasome expression by a linear regression model and the observed proteasome expression. To compute the predicted proteasome expression, the model function fit by the linear regression with the expression of *NFE2L1* as the explanatory variable and the expression of the proteasome as the response variable was used: *ŷ* = *βx* + *ε*, where *ŷ* is the predicted proteasome expression, *x* is the *NFE2L1* expression, *β* is the coefficient and *ε* is a random variable representing the noise. Then the *NFE2L1* resist score was calculated with the formula NRS = *ŷ* - *y*, where *y* is the observed proteasome expression. The three cohorts (parahippocampal gyrus from MSBB, ROSMAP and prefrontal cortex from Mayo study) were combined to compute NRS. The dataset batch effect was removed. To investigate the effect of tau aggregation on NRS, a multivariate linear regression model with the formula, *NRS ∼ Braak Score + RIN + sex + ethnicity + age + PMI*, was used. *P* < 0.05 was considered significant.

## Results

### Impaired Degradation Capacity of Proteasomes from AD Brains

A central feature of AD and other neurodegenerative disorders is the disruption of cellular proteostasis, an imbalance between protein synthesis, folding, and degradation leading to the accumulation of misfolded and aggregated proteins^29,30^. The UPS, with the 26S proteasome at its core, plays a pivotal role in maintaining neuronal proteostasis by selectively degrading aberrant or damaged proteins^4–6^. Despite its recognized importance, the precise regulation, functionality, and regional specificity of the proteasome in AD remain underexplored. Although previous studies have reported reduced proteasome activity in AD-affected brain regions^31,32^, most relied on endpoint assays of total lysates. Such assays risk capturing non-specific proteolytic activities and may not accurately reflect the intrinsic capacity of the proteasome complexes themselves. To address these limitations, we employed three complementary assays to characterize proteasome function in the neocortical Brodmann area 9 (BA9), a region heavily implicated in dementia^33,34^. First, we separated post-mortem human brain tissue into grey and white matter tissue. Grey matter represents the densely packed neuronal somata and white matter predominantly comprises myelinated axons and glial cells (oligodendrocytes, astrocytes, and microglia), this approach allowed us to determine whether proteasome function differs across these distinct cellular environments^35^.

Using kinetic fluorogenic substrate digestion over a two-hour period, proteasomes from both grey and white matter extracts in AD brains displayed a significantly reduced rate of substrate hydrolysis compared to non-cognitively impaired controls (**Fig. 1A-D**). These results suggest a widespread impairment of proteasome-mediated proteolysis in AD brains.

To ensure that these deficits were not merely reflective of altered proteasome abundance or the presence of additional proteases in crude lysates, we next purified 26S proteasomes from brain tissue using an affinity chromatography–based assay adapted from Besche and Goldberg^36^, which is a well-established method in our lab^15^. Under standardized conditions and with equal amounts of purified proteasomes, the kinetic measurements again revealed a substantially diminished degradative capacity in AD-derived 26S proteasomes compared to controls (**Fig. 1E-H**). This finding strongly supports the conclusion that intrinsic alterations within the proteasome complex, in addition to extraneous factors in the cellular milieu, underlie the observed impairment.

To further dissect the activities of distinct proteasome species, we performed in-gel activity assays under non-denaturing conditions, preserving native configurations of the 26S and 20S complexes^37^. Both 26S and 20S proteasome activities were markedly reduced in grey matter from AD brains (**Fig. 1I,J**) and although a similar trend was observed in the white matter for the 26S proteasome the 20S proteasome in white matter, did not reach statistical significance (**Fig. 1K,L**). These data suggest that proteasome subtypes may be differentially affected in a region- or cell-type-specific manner, reflecting the heterogeneity of neuronal and glial populations as well as the distinct protein quality control demands they face.

Overall, our data demonstrate that proteasomes from AD brains have a reduced capacity to degrade substrates, with persistently impaired activity even when isolated from the diseased environment. These findings imply that intrinsic and extrinsic alterations in the proteasome complex contribute to the disruption of proteostasis observed in AD.

### Proteasome Complexes are Reduced in AD Brains

Having established that proteasome activity is compromised in AD brains, we next sought to delineate how the abundance and composition of proteasome complexes are altered in AD brains. To this end, we isolated 26S proteasome-containing gel bands from in-gel activity assays and performed high-resolution proteomic analyses using PASEF -based proteomics to maximize the sensitivity and selectivity of proteomics analyses. Samples were derived from both grey and white matter of non-cognitively impaired (control) (n = 12) and AD (n = 14) brains, allowing us to evaluate potential regional differences in proteasome composition linked to AD pathology.

Quantitative proteomic and statistical analyses (using the R package DEP) identified and quantified 655 proteins in grey matter. Among the proteins, all 33–37 known proteasome subunits were present, including the alpha (PSMA1–7) and beta (PSMB1–10) subunits of the 20S core, the ATPase (PSMC1–6) and non-ATPase (PSMD1–14) subunits of the 19S regulatory particle, as well as additional sub-stoichiometric proteins associated with the 26S proteasome^38^.

A volcano plot illustrating differential analyses of 26S proteasome–enriched grey matter samples revealed 359 differentially abundant proteins, comprising 189 significantly decreased and 170 increased in AD relative to controls (**Fig.2A**). Notably, constitutive 26S proteasome subunits showed significant reductions in AD (**Fig. 2B**), indicating that the diminished proteolytic activity observed in AD (Fig. 1E–H) may stem not only from functional defects but also from reduced proteasome complex abundance. Furthermore, we detected elevated levels of aggregation-prone proteins such as tau (MAPT) and α-synuclein (SNCA) co-isolated with the proteasome, linking proteasome deficiency to the accumulation of pathological substrates on proteasomes (**Fig.2A**).

**Figure 2.**
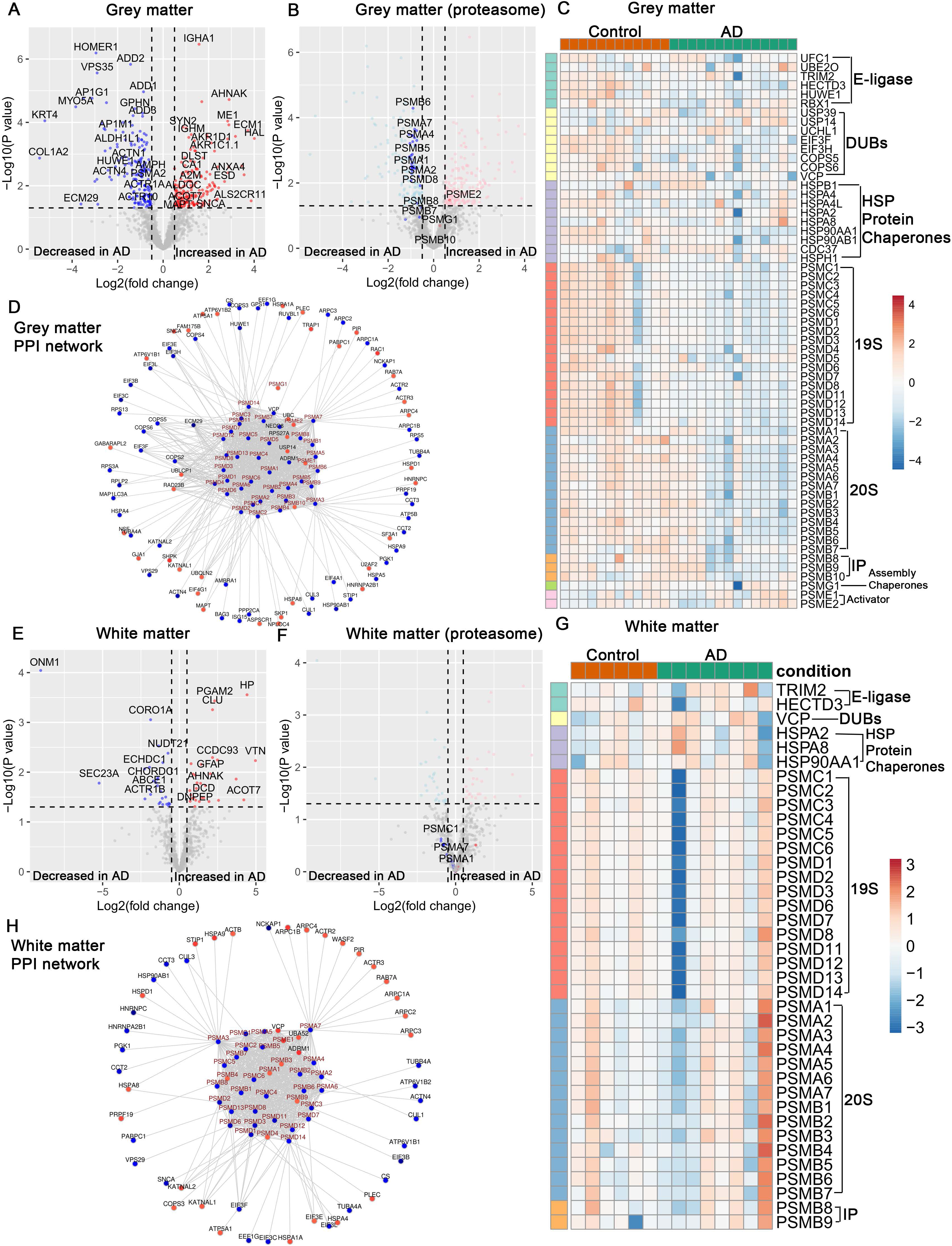
Proteomic and network analyses of gel-isolated 26S proteasomes reveal a selective depletion of proteasome subunits and associated proteostasis factors in AD grey matter compared to more moderate changes in white matter. (**A,B**) Volcano plots showing differentially abundant proteins from gel-isolated 26S proteasomes from the grey matter of AD versus control brains (control n = 12, AD n = 14). Each point represents a protein, with the x-axis indicating log2 (fold change) and the y-axis indicating –log10(p-value). The dashed vertical and horizontal lines denote significance thresholds. (**A**) Overall proteomic changes reveal numerous proteins decreased (blue) and increased (red) in AD. (**B**) Focusing on proteasome subunits and related factors, a pronounced reduction in constitutive proteasome components is observed in AD grey matter. (**C**) Heatmap of selected UPS-related proteins, including E3 ubiquitin ligases, deubiquitinating enzymes (DUBs), molecular chaperones, and proteasome subunits (19S, 20S, immunoproteasome (IP), and assembly chaperones) in grey matter. Each column represents an individual sample, and each row corresponds to a protein. The color scale indicates relative abundance changes (Z-scores), with blue representing decreased abundance and red increased abundance in AD. Grey matter shows a marked downregulation of constitutive proteasome subunits and associated assembly factors. (**D**) Protein–protein interaction (PPI) network for differentially abundant UPS-related factors in grey matter. Red nodes indicate proteins increased in AD, and blue nodes indicate proteins decreased in AD. The center of the network is dominated by reduced proteasome subunits and stabilizing factors, reflecting a core proteostatic disruption in grey matter. (**E,F**) Volcano plots for gel-isolated 26S proteasomes from white matter (n = 6, AD n = 8), analogous to panels (A,B). While significant alterations occur, the changes in proteasome subunits are less pronounced than in grey matter. Some proteins are decreased (blue) or increased (red), but constitutive proteasome components do not show as strong a depletion. (**G**) Heatmap of UPS-related proteins in white matter. Similar to (C), but the pattern is more subtle. Although some changes occur, the core proteasome subunits and assembly chaperones are not consistently downregulated, suggesting white matter proteostasis is less severely affected than grey matter. (**H**) PPI network of differentially abundant proteins in white matter. Although certain factors are altered, the network does not show the same profound depletion of proteasome subunits seen in grey matter. Some compensatory or adaptive mechanisms may help maintain proteasome functionality in white matter. Together, these analyses indicate that grey matter experiences a pronounced loss of proteasome complexes and broad destabilization of proteostasis networks in AD, while white matter remains relatively more stable, reflecting distinct region-specific vulnerabilities in the AD brain.

We extended our analysis by generating a comparative heatmap of proteasome subunits alongside proteins involved in ubiquitin-mediated proteolysis, including E3 ubiquitin ligases, deubiquitinating enzymes (DUBs), and molecular chaperones (**Fig. 2C**). While no consistent reduction was observed for upstream UPS components, we found a marked downregulation of the 19S and 20S proteasome subunits as well as immunoproteasome components in AD (**Fig. 2C**). This pattern suggests that, rather than a global UPS suppression, the proteasome complex itself is selectively vulnerable in the AD brain.

Protein–protein interaction (PPI) network analyses of first-degree proteasome interactors further supported these findings. In grey matter, proteasome subunits were consistently reduced (in blue), and several key proteostasis regulators—such as E3 ubiquitin ligases (CUL1, CUL3, HUWE1, SKP1), chaperone components (CCT2, CCT3, CCT4, CCT5, CCT6A, CCT6B, CCT7, CCT8, TCP1), HSP90-associated co-chaperones (HSP90AB1, STIP1, BAG3) and ECM29, a proteasome stabilizing factor, were also reduced (**Fig. 2D**). In contrast, tau (MAPT) and α-synuclein (SNCA) were elevated (in red), as were several molecular chaperones (HSPA1A/HSP70, HSPA8/HSC70, TRAP1, HSPD1/HSP60) and ubiquitin shuttle/DUB-related factors (UBC, RAD23B, UBQLN2, USP14, NPLOC4, UBLCP1) (**Fig. 2D**). Although these alterations likely reflect disease-stage proteostatic adjustments, the persistent loss of core proteasome subunits points to a compromised final degradation step.

In white matter, we quantified 456 proteins, including the full set of proteasome subunits. However, differential abundance analyses found only 55 proteins significantly altered (**Fig. 2E**). Unlike in grey matter, proteasome subunits in the white matter did not exhibit a statistically significant decrease (**Fig. 2F, G**). These data imply that white matter proteostasis is better preserved or more effectively compensated than grey matter, with less disruption to the proteasome complex. The PPI network revealed that although key E3 ligases (CUL1, CUL3) and select chaperones (HSP90AB1, CCT2, CCT3) were reduced (in blue) (**Fig.2H**), several chaperones, including HSP70 family members (HSPA1A, HSPA8), HSPA4, HSPA5, HSPA9, and HSPD1, as well as STIP1 (HOP), were increased (in red) (**Fig.2H**), indicative of an adaptive chaperone response.

We next performed similar quantitative proteomics and statistical analyses to identify the stochiometric and sub-stochiometric protein changes on the affinity-purified intact 26S proteasomes from both the grey and white matter of non-cognitively impaired controls and AD brains. We identified 1200 proteins in gray matter and 693 proteins in white matter. Volcano plot from gray matter of 26S proteasomes revealed 52 differentially abundant proteins, with 22 significantly decreased and 30 increased in AD relative to controls (**Fig. 3A**). For white matter affinity purified proteasome, we detected 78 differentially abundant proteins, with 43 significantly decreased and 35 increased in AD relative to controls (**Fig. 3E**). Among these changes, 26S proteasome subunits in the grey matter stood out again as a cluster of reduced proteins in AD albeit not a significant reduction. This is consistent with our finding that the mature complex of 26S proteasomes is diminished in AD (**Fig 3B**), which in turn can contribute to the reduced proteolytic capacity previously observed in this region. A heatmap of differentially abundant UPS-related proteins, including E3 ubiquitin ligases, DUBs, chaperones, and proteasome assembly factors, showed no consistent pattern. In contrast proteasome subunits were consistently downregulated (**Fig. 3C**) consistent with our results from gel-isolated proteasome showing selective vulnerability of the proteasome complex itself compared to the global UPS.

**Figure 3.**
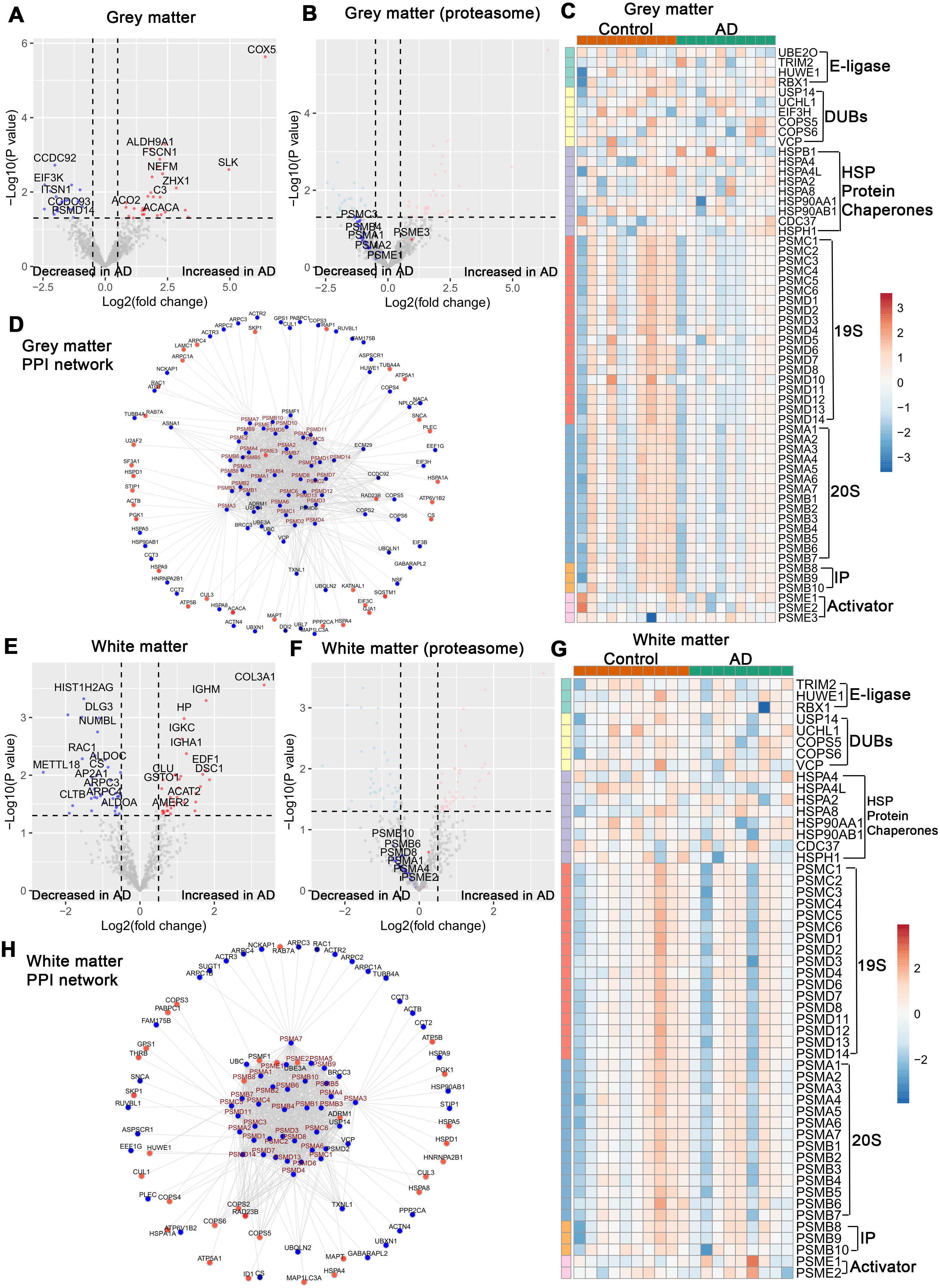
Proteomic and network analyses of affinity purified 26S proteasomes in AD. (**A,B**) Volcano plots illustrating changes in protein abundance in affinity-purified 26S proteasomes from grey matter in AD versus controls (control n = 10, AD n = 10). Each point represents an individual protein, with the x-axis showing log2 (fold change) and the y-axis displaying −log10(p-value). Dashed lines indicate significance thresholds. (**A**) Overall differentially abundant proteins are seen, with several proteins decreased (blue) or increased (red) in AD. (**B**) Focusing on proteasome subunits and related factors, a depletion of constitutive proteasome components is observed in AD grey matter even after proteasome purification. (**C**) Heatmap showing relative abundances (Z-scores) of selected UPS-related proteins in grey matter, including E3 ligases, deubiquitinating enzymes (DUBs), molecular chaperones, and proteasome subunits (19S, 20S, immunoproteasome (IP) components, and assembly factors). Each column is an individual sample, grouped by condition (Control vs. AD). The pronounced downregulation of proteasome subunits and assembly factors in AD is evident, consistent with impaired proteostasis in grey matter. (**D**) Protein–protein interaction (PPI) network of differentially abundant UPS-related factors in grey matter. Red nodes represent proteins increased in AD, while blue nodes represent decreased proteins. The network’s center is enriched for downregulated proteasome subunits and associated factors, highlighting a core proteostatic defect in AD grey matter. (**E,F**) Volcano plots of affinity purified 26S proteasome from white matter from AD and control samples (control n = 10, AD n = 9), similar to (A,B). Although some proteins are significantly altered, the depletion of proteasome components is less pronounced than in grey matter. White matter shows more subtle changes, indicating comparatively better maintenance of proteostasis. (**G**) Heatmap of UPS-related proteins in white matter, analogous to (C). While some components are altered, the pattern is less severe than in grey matter. Core proteasome subunits and chaperones are not uniformly downregulated, suggesting that proteostasis in white matter remains relatively intact or better compensated. (**H**) PPI network of differentially abundant proteins of purified proteasomes in white matter, similar to (D). Although alterations occur, the network does not reveal a profound depletion of proteasome subunits. This balanced scenario implies that white matter is less vulnerable to proteostasis collapse than grey matter. Overall, these data demonstrate a more pronounced proteasome-related proteostasis deficit in AD grey matter, while white matter is relatively spared or better adapted, underscoring region-specific differences in how AD pathology affects protein quality control systems.

The grey matter PPI network from affinity-purified samples (**Fig. 3D**) shows a dense core of proteasome subunits reduced in AD (in blue). Surrounding this core, multiple proteostasis factor including E3 ligases (CUL1, HUWE1), COP9 signalosome components (COPS2, COPS3, COPS4, COPS5, COPS6, GPS1), chaperones (HSP90AB1, CCT2, CCT3, HSPA5), and signaling factors of Nrf1 pathway such as VCP and DDI2 were also diminished (**Fig. 3D**). Moreover, key UPS adaptors and DUBs (ECM29, UBQLN1/2, USP14, and NPLOC4) showed reduced abundance as well. In contrast, certain proteins associated with purified proteasomes were increased in AD (in red), chaperones (HSPA1A, HSPA4, HSPA9), aggregation-prone AD-linked substrates such as tau/MAPT and α-synuclein/SNCA, and the ubiquitin-binding protein SQSTM1 (p62), which is known to accumulate under proteotoxic stress (**Fig. 3D**).

In white matter affinity-purified proteasome samples, we identified 693 proteins, and 78 were differentially abundant: 43 decreased and 35 increased in AD (**Fig.3E**). Although some proteasome partners were altered, the overall abundance of proteasome subunits in white matter showed no pronounced or statistically significant reductions (**Fig. 3F**). The heatmap of UPS factors in white matter revealed subtle changes without a clear pattern of proteasome depletion (**Fig. 3G**). The white matter PPI network similarly suggested a more balanced scenario: while some chaperones (CCT2, CCT3, HSPA9, HSP90, STIP1) and ubiquitin adaptors (UBC, UBEA, USP14, UBX1) were decreased (in blue), (**Fig. 3H**); the E3 ligases (CUL1, CUL3, HUWE1, SKP1), COP9 signalosome components (COPS2–6, GPS1), and certain chaperones (HSPA8, HSPD1, HSPA4, HSPA1) were increased (in red). MAPT, PSMF1, and RAD23B also rose in abundance (**Fig.3H**). Overall, these findings suggest that while white matter experiences proteostatic stress in AD, the core proteasome machinery is less severely affected than in grey matter.

In summary, our results reveal a striking regional imbalance in proteostasis in AD. Grey matter exhibits a notable depletion of proteasome complexes and a broad reduction in proteostasis factors, alongside elevated levels of aggregation-prone proteins. By contrast, white matter retains more stable proteasome levels and displays only moderate alterations. Thus, grey matter proteostasis seems more profoundly compromised, potentially driving the characteristic neurodegenerative pathology of AD in these regions. Full protein-level quantifications and supplementary data are presented in the supplementary material.

### Constitutive proteasome genes are decreased with the progression of Braak stages in AD

Our findings thus far indicate that proteasome function is diminished at both the enzymatic activity and proteomic levels in AD. To further investigate whether these impairments are reflected at the transcriptional level and to clarify how proteasome subunit gene expression relates to AD progression, we first examined bulk RNA-sequencing datasets from three large, well-characterized cohorts: the Mount Sinai Brain Bank (MSBB) with 299 brain samples from multiples regions^39^, the Religious Orders Study and Memory and Aging Project (ROSMAP) with 633 brain samples^34^ and the Mayo Clinic Study of Aging with 319 brain samples^40^. Together, these datasets encompass 1251 samples of the full spectrum of AD pathology, as indexed by Braak staging^33^. Multivariate ordinal regression models, adjusted for potential confounders (e.g., age, sex, education, ethnicity, and postmortem interval), revealed that the observed decreases in proteasome gene expression were not simply attributable to common demographic or technical variables.

Heatmaps of normalized gene expression (**Fig. 4A–C**) reveal that constitutive proteasome subunits, including both the 19S regulatory (PSMC and PSMD families) and 20S core (PSMA and PSMB families) particles, show a stark and progressive decline in expression levels starting in early Braak stages with a concomitant decrease further with increasing Braak stage (**Fig. 4A–C**). Proteasome assembly chaperones (PSMG1-4), which are critical for proper proteasome maturation and stability^38^, also exhibited a pronounced downregulation, mirroring the pattern seen in core proteasome subunits. Notably, this decline was consistently observed across all three cohorts (**Fig. 4A–C**). For MSBB dataset, this pattern of decreased proteasome gene expression was observed in multiple cortical regions (**Supplementary** Figure 5). For illustrative purposes, we present results from the parahippocampal gyrus (PHG) (**Fig.4A**), a region known for early tau pathology in AD. Supplementary analyses confirmed that other brain regions, including the frontal pole, inferior frontal gyrus, and superior temporal gyrus, exhibited similar proteasome subunit expression patterns (**Supplementary Figure 5A-C**). Interestingly, the expected decrease in proteasome expression was not observed in the cerebellum of the Mayo cohort (**Supplementary Figure 5 D**), implying regional specificity in AD-associated proteasome gene changes.

**Figure 4.**
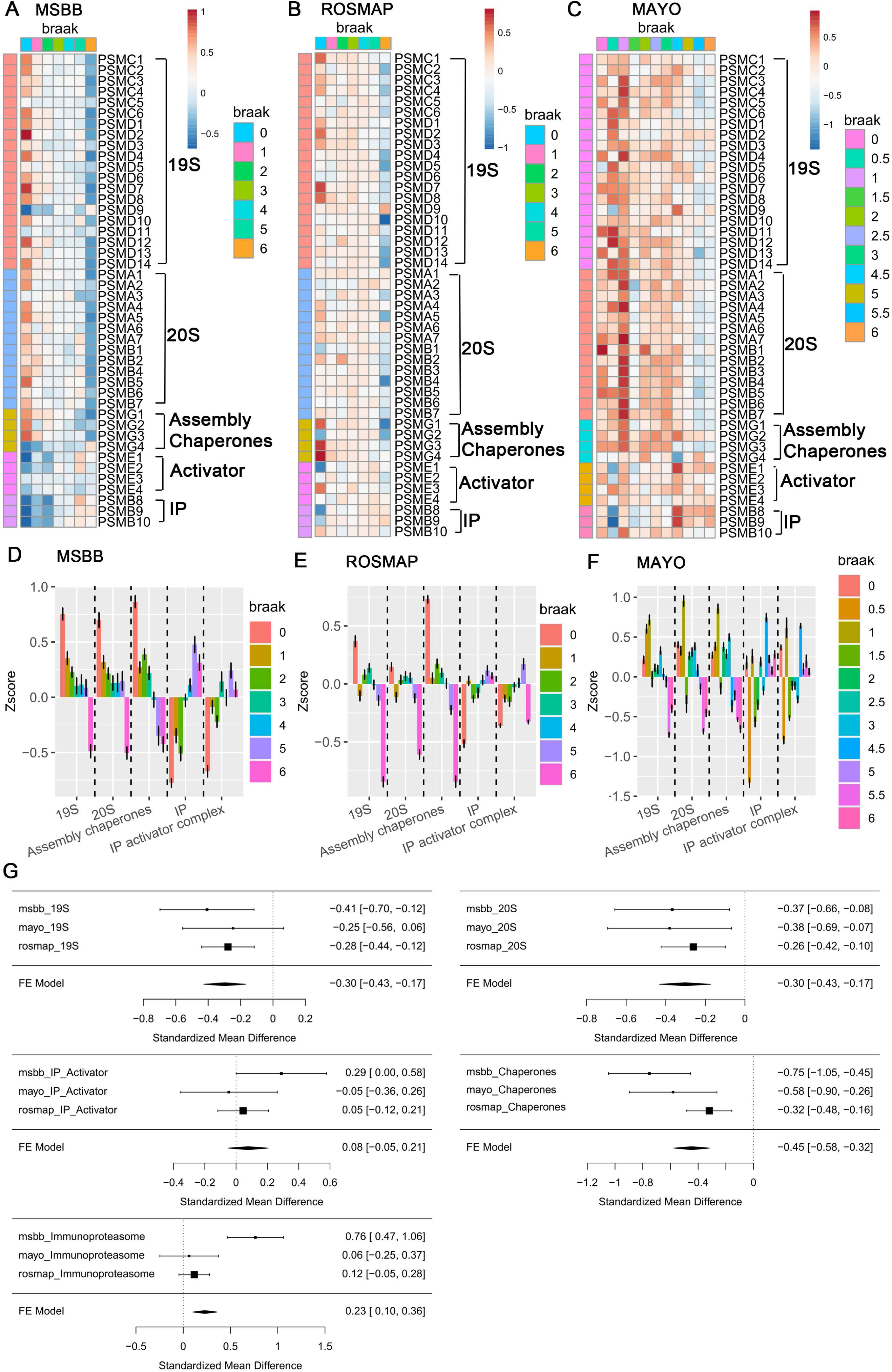
Progressive downregulation of constitutive proteasome subunits and differential responses of immunoproteasome across multiple AD cohorts. (**A–C**) Heatmaps showing the normalized expression (Z-scores) of constitutive proteasome subunits, assembly chaperones, and immunoproteasome (IP) components across increasing Braak stages for three independent cohorts of bulk-RNAseq datasets: (**A**) MSBB, (**B**) ROSMAP, and (**C**) Mayo Clinic Study of Aging. Each column represents a sample ordered by Braak stage (0-VI), while rows depict individual proteasome-related genes. Warmer colors (reds) indicate higher relative expression, and cooler colors (blues) indicate lower expression. A clear trend emerges where constitutive proteasome subunits (19S and 20S) and assembly chaperones decline with the advancing Braak stage, whereas immunoproteasome subunits show less consistent changes. (**D–F**) Boxplots of Z-scores for aggregated proteasome complexes and factors, 19S, 20S, assembly chaperones, immunoproteasome (IP), and IP activator complexes—grouped by Braak stage in (**D**) MSBB, (**E**) ROSMAP, and (**F**) Mayo cohorts. Each colored point or bar corresponds to a specific Braak stage. As Braak stage increases, 19S and 20S components consistently show a downward trend, while immunoproteasome and IP activator complexes remain relatively stable or exhibit compensatory changes. (**G**) Meta-analysis combining results from MSBB, ROSMAP, and Mayo cohorts. Forest plots present standardized mean differences across studies for key proteasome-related groups (19S, 20S, IP activator, immunoproteasome, and assembly chaperones). Negative values indicate reduced abundance in AD relative to controls. The meta-analysis confirms a robust, consistent decrease in constitutive proteasome subunits and assembly chaperones, while immunoproteasome components show more variable or modest changes. Together, these data demonstrate a reproducible and progressive loss of constitutive proteasome capacity across multiple cohorts and brain regions. The early and sustained downregulation of 19S and 20S proteasome components with AD pathology suggests a fundamental disruption in neuronal proteostasis, potentially contributing to the accumulation of aggregation-prone proteins as the disease advances.

To provide a more quantitative assessment, we aggregated proteasome genes into functionally defined complexes (19S, 20S, assembly chaperones, immunoproteasome (IP), and IP activator complexes) and plotted their Z-scores against Braak stage (**Fig. 4D–F**). The MSBB and ROSMAP datasets, which represent diverse brain regions and pathological states, consistently demonstrated significant declines in the 19S and 20S complexes as AD pathology advanced (**Fig. 4D,E**) and (**Supplementary Fig. 5 E-G**). The Mayo dataset, while slightly more variable, broadly supported this trend (**Fig. 4F**) and (**Supplementary Fig.5 H**). Collectively these results suggest that a marked impairment of the constitutive proteasome system arises early, before overt tau aggregation and worsens as tau pathology escalates.

In contrast, immunoproteasome subunits and associated immunoproteasome activator complexes did not exhibit a uniform pattern of downregulation. These components showed relative stability or an increase around Braak stage IV when constitutive proteasomes are significantly downregulated, hinting at an adaptive or compensatory response. (**Fig. 4A-F**) and (**Supplementary Fig. 5**). The immunoproteasome is more active under oxidative stress and inflammation conditions, both of which are heightened in the AD brain^41^. This divergence between the constitutive and immunoproteasome systems likely represents an alternative cellular adaptation to progressing proteotoxicity and neuroinflammation.

A fixed-effect model meta-analysis integrated data from the three cohorts and provided a robust statistical framework to summarize these findings (**Fig. 4G**). The meta-analysis confirmed the significant negative standardized mean differences for both the 19S and 20S complexes and assembly chaperones across three studies. In contrast, immunoproteasome subunits showed a smaller and less consistent effect size, underscoring their unique regulatory behavior in AD pathology (**Fig. 4G**).

Taken together, these results demonstrate a clear and reproducible pattern of declining constitutive proteasome subunit expression in the early stages of the disease and paralleling tau pathology progression in AD. Observed across multiple cohorts and brain regions, this pattern of proteasomal dysfunction underscores the importance of proteostasis failure in AD pathophysiology. Moreover, the contrasting behaviors of constitutive and immunoproteasome systems highlight the complexity of proteostatic networks and their therapeutic implications. Because proteasomal insufficiency can foster the accumulation of pathogenic proteins, strategies aimed at stabilizing or restoring proteasome function may hold promise in alleviating the neuronal damage characteristic of AD.

### snRNA-Seq Analyses Reveal Neuron-Specific Proteasome Reductions

To further dissect the cell-type–specific alterations in proteasome expression identified in our bulk RNA-seq analyses, we examined two recently published snRNA-seq datasets (Fujita et al.^27^ and Mathys et al.^28^) from dorsolateral prefrontal cortex (DLPFC) tissue samples from 619 non-overlapping brains, totaling over 3.9 million single nuclei and spanning a range of Braak stages in AD (**Fig. 5**). Building on the patterns previously observed where bulk RNA-seq analyses suggested a pronounced and progressive decline in constitutive proteasome subunits (**Fig. 4**), snRNA-seq data from ROSMAP studies reveal a neuron-specific decrease in proteasome gene expression (**Fig. 5**). These two independent datasets provide a complementary, high-resolution view of proteasome dysregulation across multiple brain cell types and across AD pathology.

**Figure 5.**
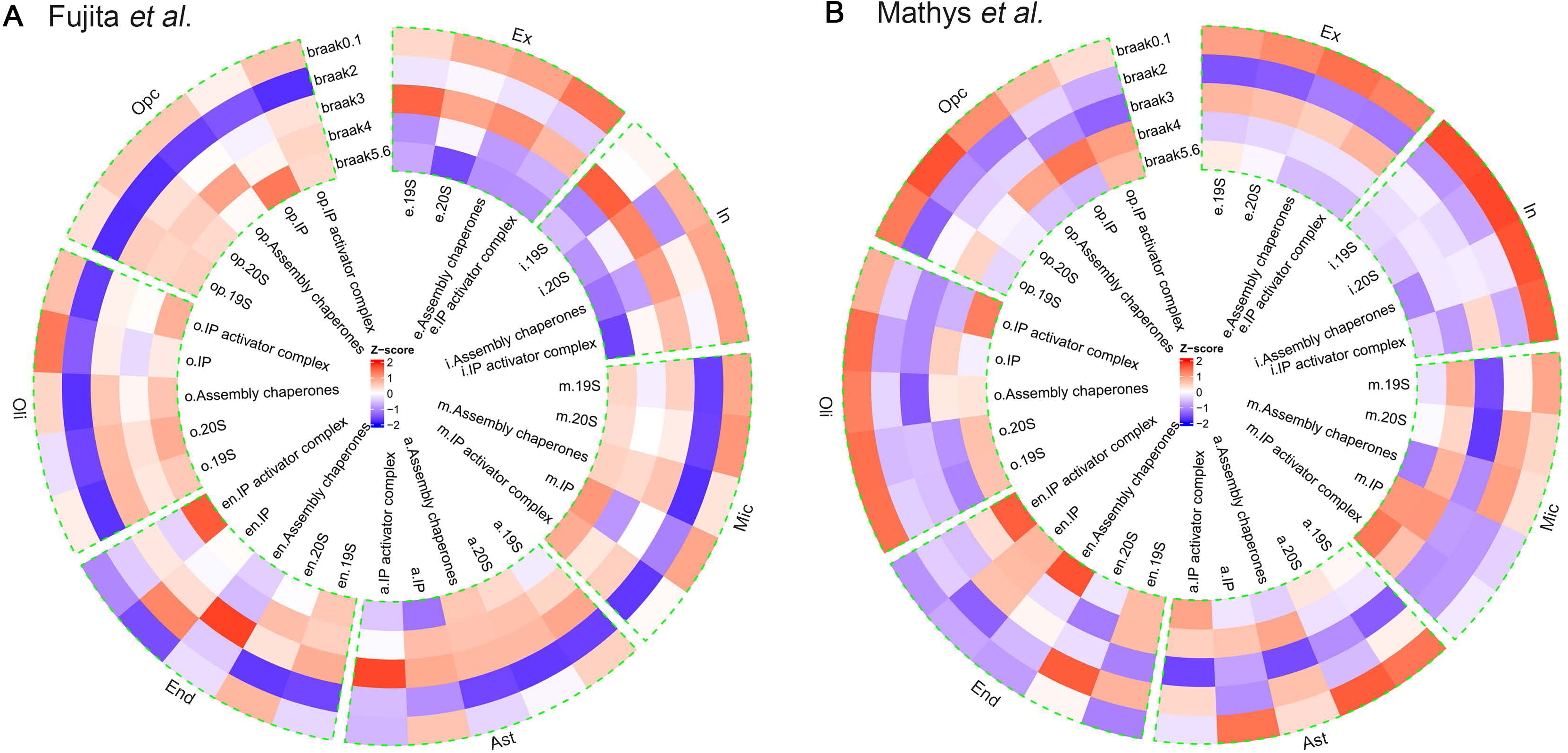
Cell-type–specific changes in proteasome-related gene expression across AD progression, as revealed by snRNA-seq datasets. (**A,B**) Circular heatmaps illustrating the Z-scored expression patterns of proteasome subunits, assembly chaperones, immunoproteasome (IP) components, and IP activator complexes across increasing Braak stages in distinct brain cell types. Each concentric ring represents a Braak stage, and each radial segment corresponds to a proteasome complex class (e.g., 19S, 20S, assembly chaperones, IP, IP activator complexes), while each wedge of the circle denotes a specific cell type. Warmer colors (reds) indicate higher relative expression, and cooler colors (blues) indicate lower relative expression. Braak stages increase outward from the center. (**A**) Data from Fujita et al.^27^ show that excitatory (Ex) and inhibitory (In) neurons progressively lose expression of constitutive proteasome subunits and assembly chaperones as Braak stage advances, shifting from red to blue hues. In contrast, non-neuronal cells (e.g., astrocytes (Ast), microglia (Mic), oligodendrocyte lineage (Oli), oligodendrocyte precursor cells (Opc), and endothelial cells (End)) maintain or even increase certain proteasome-related transcripts, indicating a more resilient or compensatory response. (**B**) Data from Mathys et al.^28^ similarly confirm a neuron-specific decline in constitutive proteasome gene expression with advancing Braak stages. Non-neuronal cell types again show relatively stable or less severely impacted proteasome-related expression patterns, consistent with the findings from Fujita et al. Together, these snRNA-seq data from two independent datasets underscore a fundamental, cell-type– dependent vulnerability in AD, with neurons showing pronounced proteasome gene downregulation as disease pathology escalates, while non-neuronal cells retain or bolster their proteostasis capacity.

In both the Fujita et al.^27^ (**Fig. 5A**) and Mathys et al.^28^ (**Fig. 5B**) datasets, concentric circular heatmaps visualize proteasome-related gene expression (19S, 20S, assembly chaperones, IP, and IP activator complexes) as a function of Braak stages, with each ring segment representing a distinct cell type and each radial segment denoting a proteasome complex class. Across increasing Braak stages, excitatory and inhibitory neurons consistently exhibit a substantial reduction in the expression of constitutive proteasome complexes and their assembly chaperones (**Fig. 5**). This neuron-specific decline aligns closely with our earlier findings from the bulk-tissue analyses of ROSMAP and other two datasets. Here extra information suggests that neurons are particularly vulnerable to proteostatic stress in AD (**Fig. 5**).

By contrast, non-neuronal cell types, including astrocytes, microglia, oligodendrocytes and endothelial cells do not follow the inferred neuronal trajectory. Instead, their expression patterns remain relatively stable or even increase for certain proteasome components, including the immunoproteasome and its activator complexes PA28αβ and PA28γ (PSME1-3) (**Fig. 5**).

These cell populations shift less dramatically across Braak stages, suggesting that they either preserve their proteasome functionality or adaptively upregulate components in response to mounting proteotoxic challenges. This differential resilience may explain the relative resistance of non-neuronal cells to pathological protein aggregation and is consistent with the more moderate changes observed in the white matter proteostasis networks reported earlier.

Taken together, these results reinforce our findings of diminished proteasome activity and subunit abundance predominantly in grey matter, aligning with the bulk RNA-seq meta-analyses underscoring a consistent downward trend in constitutive proteasome gene expression. The datasets from Fujita et al^27^. and Mathys et al.^28^ independently confirm and extend this narrative, providing robust evidence that the decline in proteasome subunit expression is not a dataset-specific artifact but a fundamental, cell-type dependent hallmark of AD progression. This convergence of evidence from multiple, independent datasets and levels of analysis strongly supports the hypothesis that neuronal proteasome insufficiency is an important contributor to the intracellular accumulation of neurotoxic proteins and the escalating proteotoxic collapse that is characteristic of AD pathology.

In contrast, non-neuronal cell types such as astrocytes, microglia, and oligodendrocytes displayed an overall increase in proteasome complex expression with advancing Braak stages. This divergence between neuronal and non-neuronal cell populations suggests that while neurons are increasingly vulnerable to protein aggregation and proteostasis collapse, non-neuronal cells may mount a more effective compensatory response, sustaining or even upregulating proteasome machinery. These findings align with the clinical and pathological observation that non-neuronal cells are generally more resistant to the hallmark protein aggregates of AD.

### Elevated NFE2L1/Nrf1 Expression Fails to Preserve Proteasome Abundance, Revealing a Regulatory Breakdown in Advanced AD

To further elucidate the molecular mechanisms underlying the progressive downregulation of proteasome transcripts in AD brains, we turned our attention to NFE2L1/Nrf1, a master transcription factor responsible for the coordinated regulation of all proteasome subunits^9,11^. To determine whether NFE2L1 directly engages proteasome gene promoters, we analyzed NFE2L1 ChIP-seq data generated by the ENCODE Project Consortium^42^ (ENCODE Project Consortium, 2012). This publicly available, high-resolution dataset derived from human cell lines enabled us to map NFE2L1 binding sites across the genome precisely.

ChIP-seq enrichment analysis revealed that proteasome-related genes exhibit significantly higher NFE2L1 occupancy than a background set of randomly selected genes, shown in the density plots (**Fig. 6A**), show that the distribution of NFE2L1 ChIP-seq signal values for proteasome genes (blue curve) is shifted toward higher enrichment compared to the background gene set (pink curve) (**Fig. 6A**). The observed difference, supported by a highly significant p-value (9.8 × 10^−4^, derived from a two-sample Kolmogorov-Smirnov test), strongly indicates that NFE2L1 preferentially binds to loci encoding proteasome subunits. These findings align with previous reports that NFE2L1 is a key transcriptional regulator of proteasome genes, functioning as an integral component of the “proteasome bounce-back” response when proteolytic capacity is challenged.

**Figure 6.**
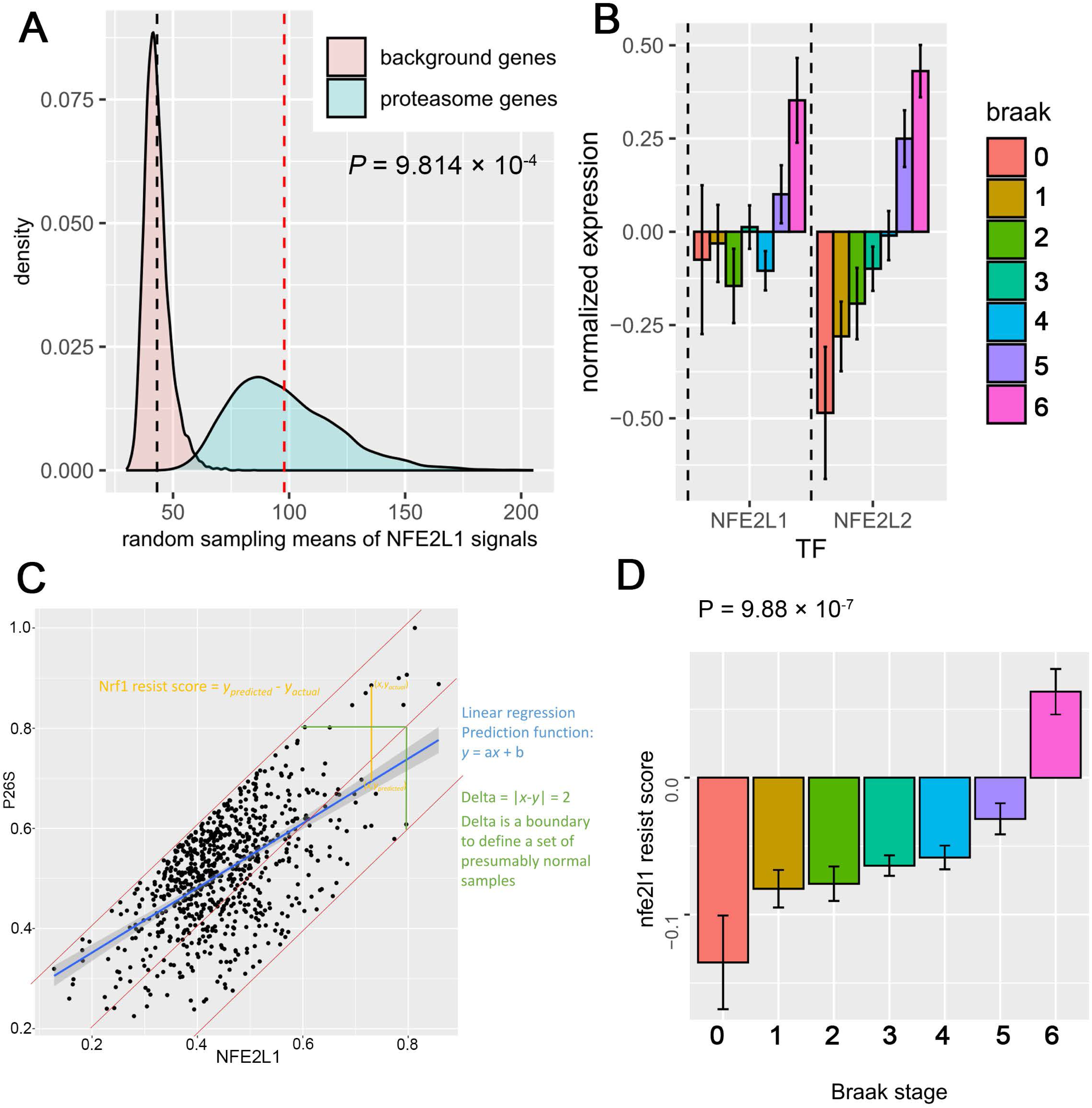
NFE2L1 preferential binding to proteasome genes but exhibits ineffective proteasome recovery in advanced AD. (**A**) Density plots of NFE2L1 ChIP-seq signal distributions for proteasome-related genes (blue) versus a background set of randomly selected genes (pink). Each curve represents the probability density of mean NFE2L1 ChIP-seq values obtained via random sampling. The vertical dashed lines indicate the median signal values for background (pink line) and proteasome (blue line) gene sets. The clear shift in the distribution toward higher values for proteasome genes, supported by a significant p-value (P = 9.814 × 10^−4^), indicates that NFE2L1 preferentially occupies proteasome gene promoters relative to random genomic loci. This enrichment highlights NFE2L1’s key regulatory role in driving proteasome gene transcription under proteostatic stress conditions. (**B**) Transcriptional changes in NFE2L1 and NFE2L2 relative to Braak stage from the three bulk RNAseq datasets (MSBB, ROSMAP and Mayo). Normalized expression data reveal that NFE2L1 and NFE2L2 transcripts increase at later Braak stages, despite declining proteasome gene expression. **(C)** The “NFE2L1 resist score” (difference between predicted vs. observed proteasome abundance based on NFE2L1 transcripts) grows more positive as Braak stage advances, indicating that elevated NFE2L1 expression does not translate into restored proteasome function (p=9.88×10^−7^). Although NFE2L1 preferentially occupies proteasome gene promoters and both NFE2L1/NFE2L2 transcripts increase at later Braak stages, these elevations fail to restore proteasome expression and function. The growing “NFE2L1 resist score” and cell-type–resolved data suggest that, despite heightened transcription factor presence, neurons cannot effectively upregulate proteasome genes as AD pathology advances. This disconnect highlights a critical failure in the compensatory mechanism intended to maintain proteostasis, contributing to the progression of Alzheimer’s disease.

To establish whether the transcriptional downregulation of proteasome subunits in AD parallels changes in NFE2L1 expression, we analyzed bulk RNA-seq data from the three independent cohorts across increasing Braak stages of AD pathology. We also examined NFE2L2 (Nrf2), a closely related transcription factor reported to induce several proteasome genes under oxidative stress conditions^43,44^. Surprisingly, our results showed that while proteasome gene expression declined as Braak staging advanced, both NFE2L1 and NFE2L2 mRNA levels actually increased at later Braak stages (**Fig. 6B**). This divergence suggests that even though the transcriptional regulators are upregulated or maintained, the proteasome transcription fails to be restored. Potential reasons for this disconnect could include impaired NFE2L1 transcriptional activity, and impaired ER to nucleus pathway which could dampen the induction of proteasomes.

To quantitatively link NFE2L1 expression with proteasome expression, we applied a linear regression model relating NFE2L1 transcript levels to measured proteasome 26S (P26S) content (**Fig. 6C**). Using the model’s predicted P26S values (y_predicted) based on NFE2L1 expression to the actual observed values (y_actual), we derived an “NFE2L1 resist score” defined as (y_predicted – y_actual). This resist score characterizes how closely a sample’s proteasome expression aligns with what would be expected from its NFE2L1 expression. Samples whose P26S levels are in line with NFE2L1-driven predictions remain near zero, whereas those with lower-than-expected P26S content (positive scores) indicate impaired proteasome transcription despite adequate transcriptional input, or as the scoring name implies, “resist” to the transcription of NFE2L1, and those with higher-than-expected P26S transcripts (negative scores) may reflect compensatory or adaptive mechanisms maintaining proteostasis. Setting a boundary (**Δ** = 2) allowed us to identify samples considered “normal” with regard to NFE2L1–proteasome coupling. Samples outside this boundary deviate substantially from the predicted relationship (**Fig. 6C**). Notably, when we examined this score by Braak stage, we observed that early-stage samples, Braak 0, are below 0 (negative scores), indicating a healthy proteasome content and activity with high proteasome abundance and low NFE2L1 expression (**Fig. 6D**) However, individuals with greater AD pathology the NFE2L1 resist score increased significantly (P = 9.88 × 10^-7^), becoming distinctly positive in individuals at Braak stages V and VI (**Fig. 6D**). In these advanced stages, higher NFE2L1 transcript levels did not translate into comparable increases in proteasome abundance, suggesting a breakdown in the transcription-to-proteasome assembly axis.

This finding implies that the cell signaling, such as the Nrf1-bound ER to nucleus cascade, progressively fails in the AD brain. Suggesting that, while NFE2L1 indued transcription may attempt to compensate for proteostatic stress in AD and the transcriptional activity of the Nrf1 pathway necessary for proteasome gene expressions becomes impaired. These results underscore the complexity of proteostasis regulation in AD, in which elevated transcriptional regulators alone are insufficient to maintain proteolytic capacity. Understanding the mechanisms governing Nrf1 activity and proteasome induction will be crucial for restoring proteostasis and mitigating neuronal damage in AD.

### Nrf1 Stabilization and Impaired Proteasome “Bounce-Back” Response in AD Brains

To assess the transcriptional regulatory capacity of Nrf1 in AD, we first examined its protein abundance and localization in proteasome-enriched (soluble) cortical (BA9 region) extracts from both control and AD brains. Western blot analyses revealed that total Nrf1 protein, the lower and upper bands, were significantly increased in AD samples (**Fig. 7A,B**). In contrast, PSMG1, a proteasome assembly chaperone essential for proper proteasome maturation, was markedly reduced (**Fig. 7A,C**). Rpt5/PSMC3, a subunit of the 19S regulatory particle, showed a nonsignificant trend toward reduction (**Fig. 7A,D**). The stabilization of Nrf1 in AD suggests that its expected proteasome-mediated degradation is reduced, likely reflecting diminished proteasome activity. Under normal conditions, robust proteasome activity continuously degrades Nrf1, maintaining its low basal levels. In AD, however, impaired proteasome function slows Nrf1 turnover, allowing it to accumulate. This process parallels the well-characterized “bounce-back” response, a cellular mechanism wherein reduced proteasome activity leads to the stabilization of Nrf1 and translocation of Nrf1 to the nucleus, which in turn should transcriptionally upregulate proteasome subunit genes to restore proteolytic capacity. Yet, despite elevated Nrf1, AD brains fail to mount the expected compensatory increase in proteasome subunits. Instead, reduced PSMG1 and Rpt5/PSMC3 suggest that while Nrf1 levels rise, the downstream signaling steps required to enhance proteasome content do not occur effectively, indicating a stalled bounce-back response.

**Figure 7.**
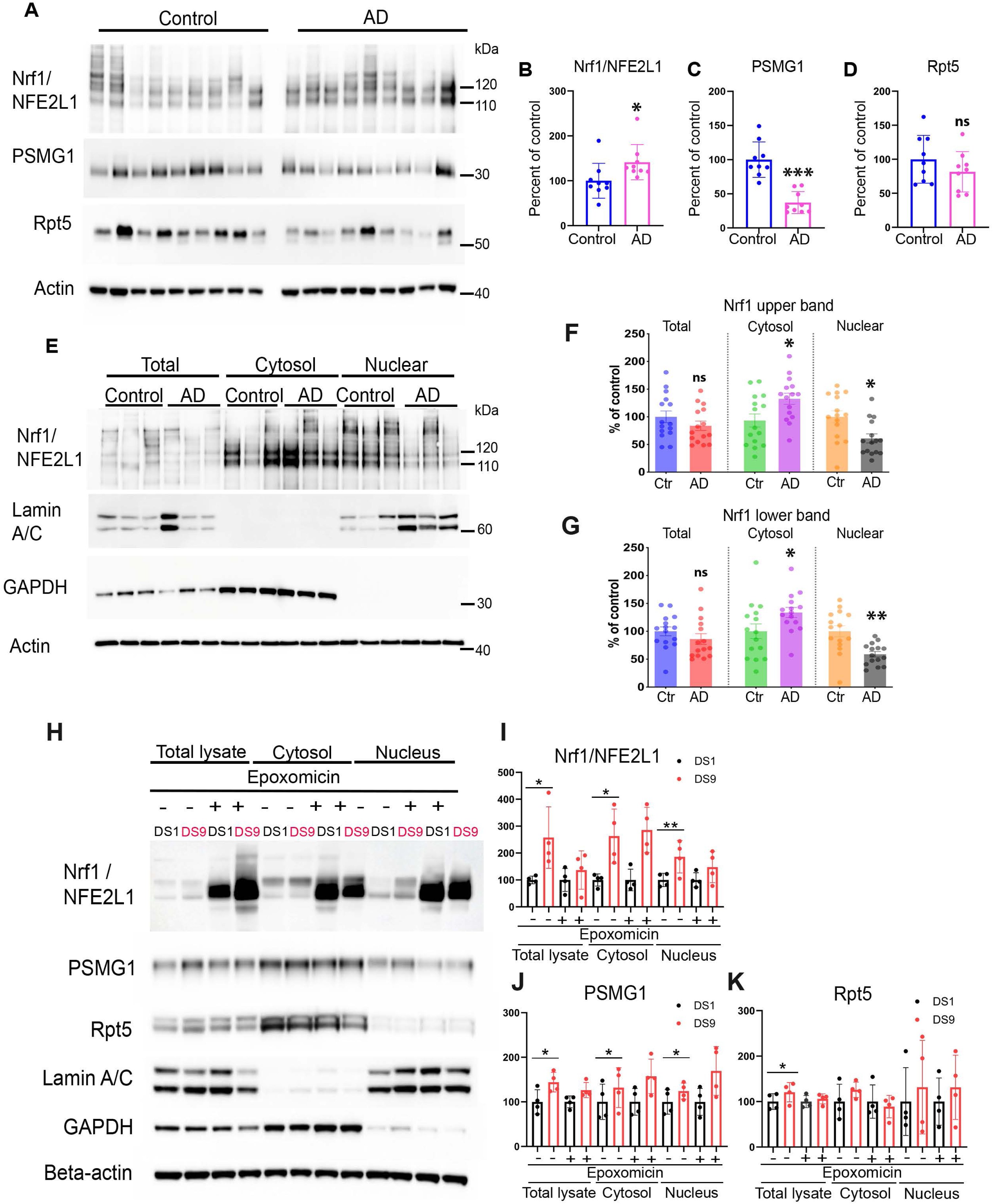
Impaired Nrf1 nuclear translocation in AD and an impaired bounce-back response of proteasomes. (**A–D**) Western blot analysis of Nrf1, PSMG1 and Rpt5/PSMC3 in proteasome-enriched soluble cortical extracts from control and AD brains (control n = 9, AD n = 9). (**A**) Representative immunoblots. Actin serves as a loading control. (**B–D**) Quantifications show that (**B**) total Nrf1 levels are significantly elevated in AD, while (**C**) PSMG1 is markedly reduced and (**D**) Rpt5/PSMC3 levels are not significantly reduced. (**E–G**) Subcellular fractionation of cortical extracts to assess Nrf1 localization. (**E**) Representative blots of total, cytosolic, and nuclear fractions from control and AD brains (control n = 15, AD n = 15). Lamin A/C and GAPDH are used as nuclear and cytosolic markers, respectively, and actin as a loading control. (**F,G**) Quantifications of upper and lower Nrf1 bands (corresponding to different post-translationally modified forms) show that total Nrf1 levels were unchanged between groups whereas the cytosolic levels were increased in AD. Contrary to the cytosolic Nrf1, nuclear Nrf1 is significantly decreased in AD. This suggests impaired nuclear translocation or processing of Nrf1 required for effective transcriptional activation of proteasome genes. **(I)** Representative Western blots showing Nrf1, PSMG1 and Rpt5 in total lysates, cytosolic, and nuclear fractions of two cell lines (DS1 and DS9) treated with or without epoxomicin, a proteasome inhibitor (four biological experiments). Lamin A/C serves as a nuclear marker, GAPDH as a cytosolic marker, and β-actin as a loading control. **(B–D)** Quantifications of (**B**) Nrf1, (**C**) PSMG1, and (**D**) Rpt5 levels comparing DS9 to DS1, a control condition. In the absence of epoxomicin, Nrf1 (upper and lower bands) undergoes rapid degradation, maintaining low basal levels (DS1cells, control condition). Upon reduced proteasome activity under persistent tau aggregation (DS9 cells condition) Nrf1 upper and lower bands increase in all the fractions. Upon proteasome inhibition with epoxomicin, cytosolic Nrf1 accumulate in the nucleus, indicative of the activated “bounce-back” response aimed at restoring proteasome capacity. This response includes upregulation of PSMG1 in both total and nuclear fractions. Rpt5 levels also show modest changes. These results demonstrate that pharmacological proteasome inhibition can recapitulate aspects of the compensatory mechanism attempting to restore proteasome homeostasis and highlight the enhanced responsiveness in a proteostasis-compromised cell line (DS9). Data are presented as mean ± SEM; each point represents an individual sample; ns = not significant, *p<0.05, **p<0.01, ***p<0.001.

Subcellular fractionation experiments provided additional insight into Nrf1 dysregulation (**Fig. 7E–G**). Immunoblotting of cytosolic and nuclear fractions showed that AD samples contain more cytosolic Nrf1 and significantly less nuclear Nrf1 compared to controls (**Fig. 7E–G**). These data suggest that although Nrf1 accumulates in AD, it is not efficiently translocated to the nucleus. As a result, Nrf1 cannot fully execute its transcriptional activity to compensate for reduced proteasome activity, underscoring a failure of the protective bounce-back mechanism in the diseased brain. The tissue localization control proteins such as Lamin A/C (nuclear marker), GAPDH, and β-actin (cytosolic/loading controls) confirmed the successful fractionation of cytosolic and nuclear proteins (**Fig. 7E**).

### Cellular Models Confirm Nrf1-Mediated Proteasome Bounce-Back Response Under Proteasome Impairment

To further investigate how impaired or reduced proteasome activity affects the bounce-back response, we examined two cell lines, DS1 and DS9 clones generated from HEK 293 cells^25^ which both express truncated diseases associated with mutant tau but differ in their proteostasis status. DS1 cells model relatively intact proteasome function since they don’t exhibit tau aggregation, whereas DS9 cells, with persistent tau aggregation, mimic a compromised proteasome state. Both cell lines were treated with 50nM epoxomicin overnight, a potent proteasome inhibitor. In DS9 cells, western blot analyses revealed significantly increased Nrf1 levels in the total, cytosolic and nuclear fraction compared to DS1 cells (**Fig. 7 H,I**), consistent with reduced proteasome-mediated degradation. Importantly, this increase was accompanied by elevated PSMG1 and Rpt5/PSMC3 levels (**Fig. 7 H,J,K**), indicating that under reduced proteasome activity due to tau aggregation, the expected bounce-back response, including Nrf1 stabilization, nuclear translocation and subsequent proteasome component upregulation was engaged in DS9 cells (**Fig. 7 H,J,K**). The complete nuclear accumulation of Nrf1 upon epoxomicin treatment confirmed that inhibiting proteasome activity triggers massive Nrf1 translocation to the nucleus, driving proteasome gene expression to restore proteolytic capacity (**Fig. 7 H,I**). However, toxicity associated with epoxomicin exposure may limit the full effectiveness of this compensatory mechanism. The localization control proteins such as Lamin A/C (nuclear marker), GAPDH, and β-actin (cytosolic/loading controls) confirmed the successful fractionation of cytosolic and nuclear proteins (**Fig. 7H**).

Overall, our findings demonstrate that while Nrf1 stabilization and nuclear translocation typically support a crucial compensatory response to diminishing proteasome activity, this system is disrupted in AD. In our cellular models, Nrf1 responds predictably to proteasome inhibition by moving to the nucleus and increasing proteasome levels; however, in AD, these subsequent processes seem impaired. This disconnect highlights that the pathological brain cannot effectively utilize a normally protective mechanism, ultimately contributing to proteostatic failure and the progression of AD.

## Discussion

Our integrated findings reveal a comprehensive portrait of proteasome insufficiency in AD that spans across multiple levels of regulation. From the reduced proteasome gene expression in the early stages of the disease to the eventual collapse of proteolytic capacity and a stalled transcriptional “bounce-back” response, our data collectively emphasize that proteasome dysfunction should be viewed as a core feature of AD pathogenesis rather than merely a downstream consequence of widespread protein aggregation. One of the most striking observations is that the transcripts for constitutive proteasome subunit decline during the early Braak stages, before overt tau aggregation and neurofibrillary tangle formation are fully established. This early transcriptional decline indicates that proteasome deficits emerge as an early event in the disease course, predisposing neurons to accelerated protein aggregation and neuronal injury. As tau pathology and other proteotoxic stresses increase, the transcriptional suppression of proteasome genes becomes more pronounced, reinforcing a vicious cycle in which diminished proteasomes promote further protein misfolding and accumulation. Consistent with these transcriptional changes, our enzymatic assays demonstrate that proteasome function is impaired in AD. Even after purification to remove extraneous cellular components, AD-derived proteasomes show an intrinsically reduced ability to degrade substrates. This finding suggests an additional structural or compositional alteration within the proteasome itself, beyond extrinsic factors contributing to the observed functional decline.

Quantitative proteomics builds upon these functional and transcriptional deficits by revealing a widespread reorganization of proteostasis-related networks. The significant reduction of constitutive proteasome subunits in grey matter, particularly in neuron-rich regions that are severely affected, coincides with lower levels of proteasome assembly chaperones. Accompanying this impairment of the proteolytic function is the accumulation of aggregation-prone substrates, such as tau, α-synuclein and p62, linking proteasome insufficiency directly to the pathogenic protein aggregates, which can in turn further inhibit proteasome activity. In contrast, white matter shows a more moderate response, with proteasome subunits. This regional and cellular specificity further supported by snRNA-seq data highlights that non-neuronal cells can maintain or adapt their proteostasis mechanisms more effectively than neurons. As a result, neuronal populations experience a more precipitous loss of proteolytic capacity, aligning with the early transcriptional deficits and creating a permissive environment for AD pathology to progress. A key unresolved issue is why intrinsic compensatory pathways fail to mitigate these early proteasome deficits. Under normal conditions, reduced proteasome activity stabilizes Nrf1, a master transcription factor that translocates to the nucleus to induce proteasome gene expression, known as the “bounce-back” response. Our ChIP-seq results confirm that NFE2L1 is directly associated with proteasome gene promoters, and the elevated levels of NFE2L1 transcripts we observe imply an attempt at compensation as AD progresses. Despite this, the expected transcriptional reactivation of proteasome subunits does not take place. Instead, the Nrf1 protein accumulates in the cytosol in AD brains, failing to effectively enter the nucleus and reactivate proteolytic capacity. This disconnect between elevated Nrf1 expression and inadequate proteasome gene induction highlights a breakdown in the signaling cascade essential for executing the bounce-back response.

Hence, early transcriptional downregulation of proteasome genes and impaired proteasome activity, combined with impaired Nrf1 nuclear translocation, suggest that neurons lose both their baseline proteostatic stability and their ability to compensate as the disease progresses. Rather than preemptively reinforcing proteasome capacity in response to unfolding proteotoxic challenges, AD neurons become locked into a downward spiral of reduced proteasome availability and escalating protein aggregation. Our data thus propose that proteostasis dysfunction is not simply downstream of tau and amyloid pathology but represents a critical upstream factor that renders neurons vulnerable to harmful protein accumulations. By demonstrating that proteasome genes are suppressed before overt tau pathology emerges, we identify proteasome deficits as one of the possible initiators of later pathogenic processes. From a therapeutic standpoint, these findings argue for early interventions to enhance proteasome function or stabilize Nrf1-mediated transcriptional responses.

In summary, our integrated analyses show that proteasome dysfunction in AD is multifaceted, emerging early and progressively worsening as the disease advances. The initial transcriptional downregulation of proteasome genes sets the stage for persistent proteasome depletion, reduced enzymatic activity, and ineffective compensatory responses, particularly involving Nrf1. These insights emphasize proteasome insufficiency as a central, upstream driver of AD pathology, presenting new opportunities for early interventions that restore proteostasis and potentially delay or prevent the full onset of the disease.

## Data availability

All data supporting the findings of this study are provided within the main text and supplementary materials, including figures and corresponding Excel files. Datasets from bulk RNA-seq and snRNA-seq are available from AD Knowledge Portal: https://www.synapse.org/Synapse:syn2580853/wiki/409840, and NFE2L1 ChIP-seq data is available from ENCODE: https://www.encodeproject.org/experiments/ENCSR543SBE/.

## Acknowledgments

We thank Dr. Marc Diamond for providing us with the HEK-293 RD-YFP (DS1 and DS9 clone) cell lines. We thank the Columbia University Alzheimer’s Disease Research Center (ADRC), funded by NIH grant P30AG066462 to S.A. Small (P.I.) for providing recourses for the study. We thank Dr. Andrew Teich, the Director of The New York Brain Bank (NYBB) at Columbia University, for providing us with postmortem brains.

## Funding

This work was funded by the NIA R01AG070075 and R01AG064244, awarded to N.M

## Competing interests

The authors report no competing interests.

## Supplementary material

Supplementary material is available at *Brain* online

## Notes

### Competing Interest Statement

The authors have declared no competing interest.

