## Supplementary material for "Early Proteasome Gene Downregulation And Impaired Proteasomes Function Underlie Proteostasis Failure In Alzheimer’s Disease"

**TABLE 1: Human demographics and neuropathological data**.

| Case # | Sex | Age at Death | PM-C | PMI-F | NPDX |
|---|---|---|---|---|---|
| 5732 | F | 82 | 3:30 | 18:28 | ADNC |
| 278 | M | 86 | 2:05 | 50:25 | ADNC |
| 250 | F | 89+ | 2:15 | 6:16 | ADNC |
| 279 | M | 83 | 3:45 | 5:35 | ADNC |
| 218 | F | 86 | 5:10 | 6:20 | ADNC |
| 5667 | F | 89 | 3:00 | 29:25 | ADNC |
| 5650 | M | 89 | 3:30 | 9:55 | ADNC |
| 5675 | F | 80 | 2:45 | 38:25 | ADNC |
| 5671 | M | 78 | 1:15 | 26:30 | ADNC |
| 5694 | F | 79 | 2:00 | 10:16 | ADNC |
| 5753 | F | 89+ | | 34:35 | ADNC |
| 5729 | F | 76 | | 20:00 | ADNC |
| 5708 | F | 89+ | | | ADNC |
| 5706 | F | 80 | | 11:57 | ADNC |
| 5779 | F | 78 | | | ADNC |
| 158 | M | 71 | 22:15 | 62:20 | ADNC |
| 200 | M | 83 | 10:30 | 11:40 | ADNC |
| 239 | M | 86 | 2:43 | 4:03 | ADNC |
| 255 | F | 81 | 10:45 | 15:00 | ADNC |
| 323 | F | 89 | 4:40 | 6:40 | ADNC |
| 324 | F | 85 | 4:15 | 8:35 | ADNC |
| 344 | F | 89 | 5:00 | 7:30 | ADNC |
| 352 | F | 89 | 3:10 | 14:05 | ADNC |
| 4069 | M | 87 | 4:24 | 8:25 | ADNC |
| 1371 | F | 66 | 3:15 | 14:23 | ADNC |
| 2013 | M | 84 | 18:21 | 20:56 | ADNC |
| 3865 | F | 89+ | 19:22 | 21:17 | ADNC |
| 4099 | M | 69 | 3:55 | 16:35 | ADNC |
| 4103 | F | 87 | 5:45 | 6:55 | ADNC |
| 4135 | M | 75 | 4:55 | 6:10 | ADNC |
| 4160 | F | 88 | 5:35 | 6:45 | ADNC |
| 4310 | F | 86 | 0:10 | 4:50 | ADNC |
| 4454 | M | 88 | 2:00 | 16:55 | ADNC |
| 4513 | M | 84 | | 7:10 | ADNC |
| 4865 | M | 89+ | 3:45 | 8:35 | ADNC |
| 4857 | M | 89+ | | 12:40 | ADNC |
| 5133 | M | 87 | | 5:15 | ADNC |
| 5359 | F | 89+ | 11:00 | 22:50 | ADNC |
| 5420 | F | 89+ | 1:25 | 6:02 | ADNC |
| 4033 | F | 88 | 14:02 | 15:22 | ADNC |
| 2467 | M | 89 | | 6:15 | ADNC |
| 4448 | M | 89 | | 12:50 | ADNC |
| 4341 | M | 83 | | 15:00 | ADNC |

| | | | | | |
|---|---|---|---|---|---|
| 5296 | M | 67 | | 8:25 | ADNC |
| 5365 | M | 87 | | 6:00 | ADNC |
| 5367 | M | 73 | | 23:00 | ADNC |
| 5398 | F | 89 | 1:00 | 19:25 | ADNC |
| 5556 | M | 70 | 0:30 | 8:37 | ADNC |
| 5333 | M | 89 | 0:30 | 7:17 | ADNC |
| 5475 | F | 68 | 0:30 | 9:20 | ADNC |
| 5481 | M | 83 | | 17:16 | ADNC |
| 5510 | M | 89+ | 7:40 | 19:21 | ADNC |
| 5453 | F | 89+ | 2:15 | 6:00 | ADNC |
| 5521 | M | 67 | | 14:55 | ADNC |
| 5700 | M | 75 | | 19:20 | ADNC |
| 5749 | F | 89 | | 22:55 | ADNC |
| 782 | M | 83 | 5:45 | 10:15 | ADNC |
| 4096 | F | 87 | 4:24 | 8:25 | ADNC |
| 5708 | M | 54 | 2:40 | 17:02 | ADNC |
| 187 | F | 80 | 4:20 | 7:40 | CTR |
| 2465 | F | 83 | 6:50 | 9:15 | CTR |
| 2791 | F | 80 | 4:30 | 6:20 | CTR |
| 5405 | F | 80 | 13:05 | 20:05 | CTR |
| 5420 | F | 80 | 5:40 | 11:40 | CTR |
| 4590 | F | 79 | 2:10 | 12:30 | CTR |
| 328 | M | 89+ | 4:47 | 11:17 | CTR |
| 4070 | F | 89+ | 5:25 | 10:40 | CTR |
| 5382 | M | 62 | | 5:24 | CTR |
| 159 | F | 69 | 3:40 | 15:40 | CTR |
| 187 | F | 80 | 4:20 | 7:40 | CTR |
| 360 | M | 74 | 5:15 | 23:45 | CTR |
| 2465 | F | 83 | 6:50 | 9:15 | CTR |
| 2791 | F | 89+ | 4:30 | 6:20 | CTR |
| 4105 | F | 89+ | 13:05 | 20:05 | CTR |
| 5133 | M | 87 | | 5:15 | CTR |
| 5181 | M | 89+ | | 7:58 | CTR |
| 5385 | M | 88 | 2:45 | 14:03 | CTR |
| 5420 | F | 89+ | 1:25 | 6:02 | CTR |
| 5514 | F | 89+ | | 9:24 | CTR |
| 199 | F | 82 | 3:50 | 22:45 | CTR |
| 346 | M | 84 | 10:00 | 14:10 | CTR |
| 5404 | F | 54 | 6:41 | 16:36 | CTR |
| 5696 | M | 84 | | 29:10 | CTR |
| 242 | M | 67 | 4:43 | 7:18 | CTR |
| 4523 | F | 62 | 4:33 | 22:38 | CTR |
| 5227 | M | 75 | | 7:35 | CTR |
| 5518 | F | 89+ | | 11:35 | CTR |
| 4915 | M | 62 | | 40:05 | CTR |
| 4931 | M | 66 | | 25:00 | CTR |
| 5382 | M | 62 | | 5:04 | CTR |
| 5404 | F | 52 | | 16:40 | CTR |

| | | | | | |
|---|---|---|---|---|---|
| 319 | F | 58 | 2:09 | 25:34 | CTR |
| 5686 | M | 84 | | 15:10 | CTR |
| 276 | M | 71 | | 12:40 | CTR |
| 171 | M | 70 | | 5:32 | CTR |
| 147 | F | 63 | | 24:20 | CTR |
| 305 | M | 80 | | 17:13 | CTR |
| 108 | M | 68 | | 9:00 | CTR |

*PMI-C, post-mortem interval – time to cold (h:min); PMI-F, post-mortem interval – time to frozen (h:min); NPDX, Neuropathologic diagnosis; ADNC, Alzheimer's Disease Neuropathologic Changes. CTR, Control.*

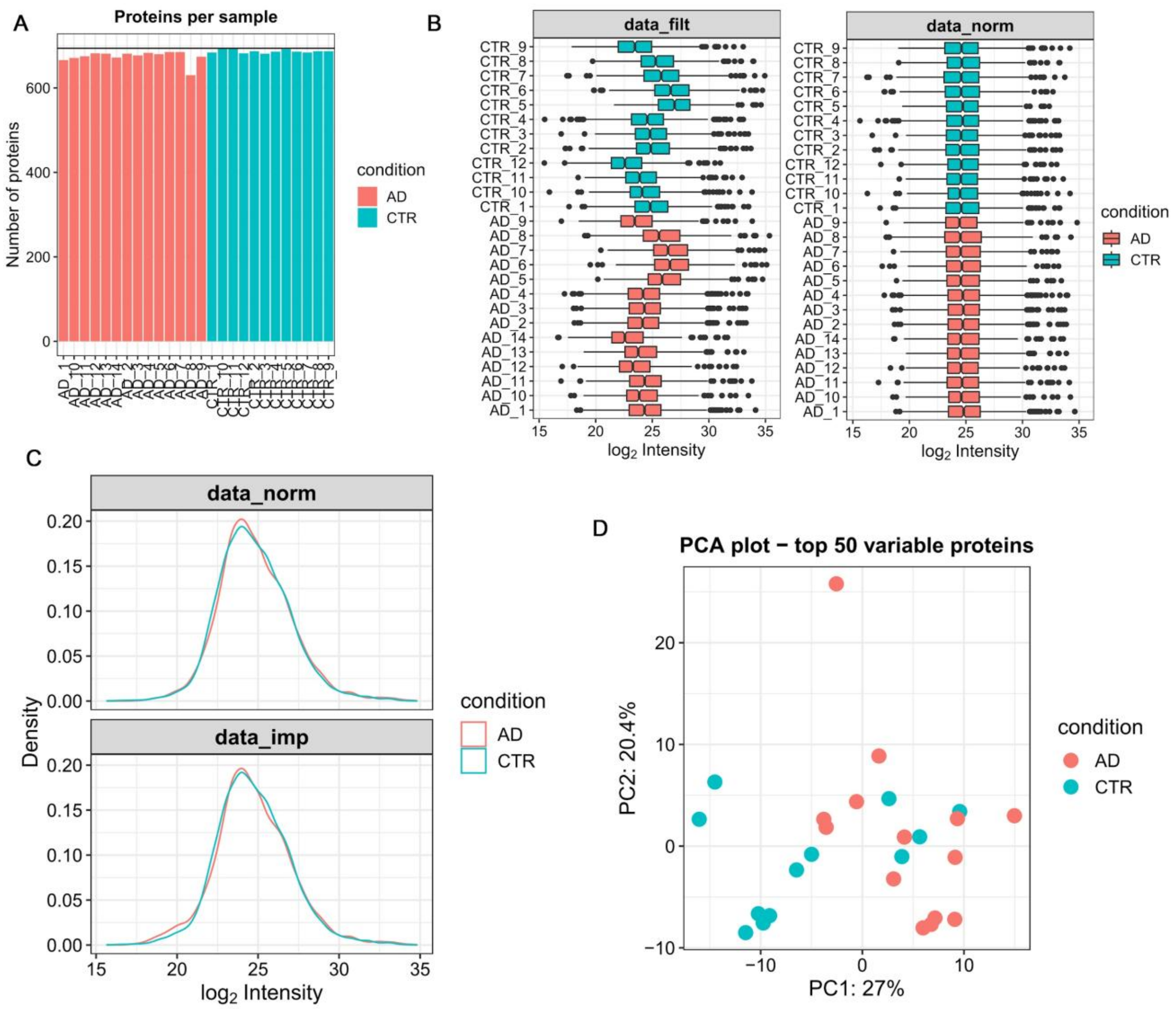


**Supplementary Figure 1.**

**Preprocessing for gel-isolated proteasome proteomics from grey matter.**

(**A**) Bar plot displaying the number of proteins identified in each individual sample. Stringent filtering removed proteins not detected in all replicates of at least one condition (control or AD). (**B**) Box plots comparing the distributions of log-transformed peak intensities for each sample before and after variance-stabilizing normalization (VSN). This panel highlights how VSN reduces technical variation across samples. (**C**) Density plots illustrating the distributions of log-transformed peak intensities before and after missing data imputation via a left-shifted "MinProb" distribution. The imputation step helps approximate low-abundance signals not captured in the raw data. (**D**) Principal component analysis (PCA) scatter plot using the top 50 most variably expressed proteins. The plot is used to visualize potential batch effects and examine overall data structure between control and AD samples

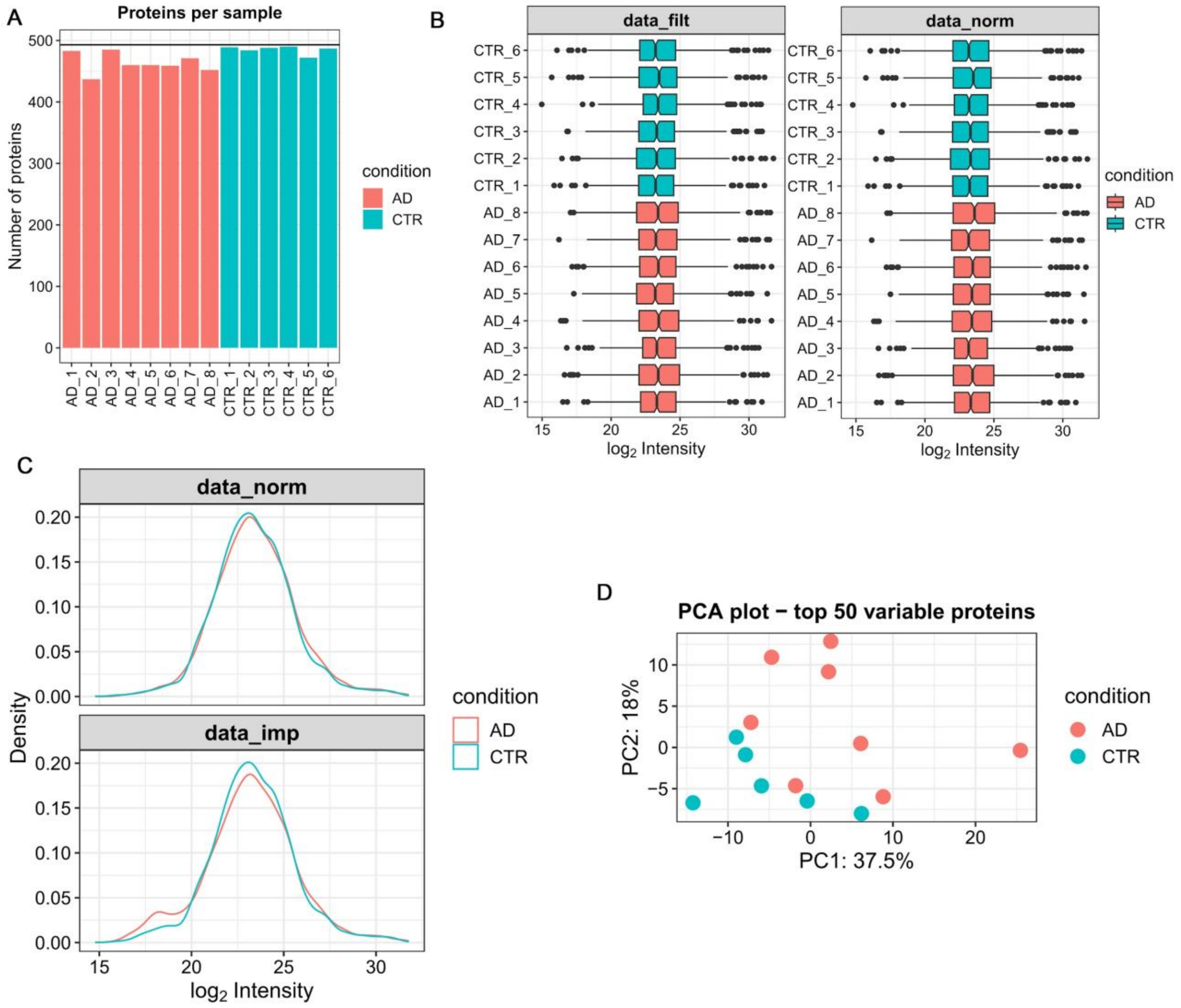


**Supplementary Figure 2**

**Preprocessing Workflow for Gel-Isolated Proteasome Proteomics from White Matter**

(**A**) Bar plot illustrating the number of proteins detected in each individual sample. Proteins not identified in all replicates of at least one condition (control or AD) were removed prior to downstream analyses. (**B**) Box plots showing the distributions of log-transformed peak intensities across samples before and after variance-stabilizing normalization (VSN). This panel demonstrates how VSN reduces technical variation. (**C**) Density plots comparing the distributions of log-transformed peak intensities prior to and following missing data imputation using a left-shifted “MinProb” distribution. Imputation compensates for low-abundance signals absent from the raw dataset. (**D**) Principal component analysis (PCA) scatter plot using the top 50 most variably expressed proteins. The PCA highlights overall data structure and potential batch effects between control and AD samples.

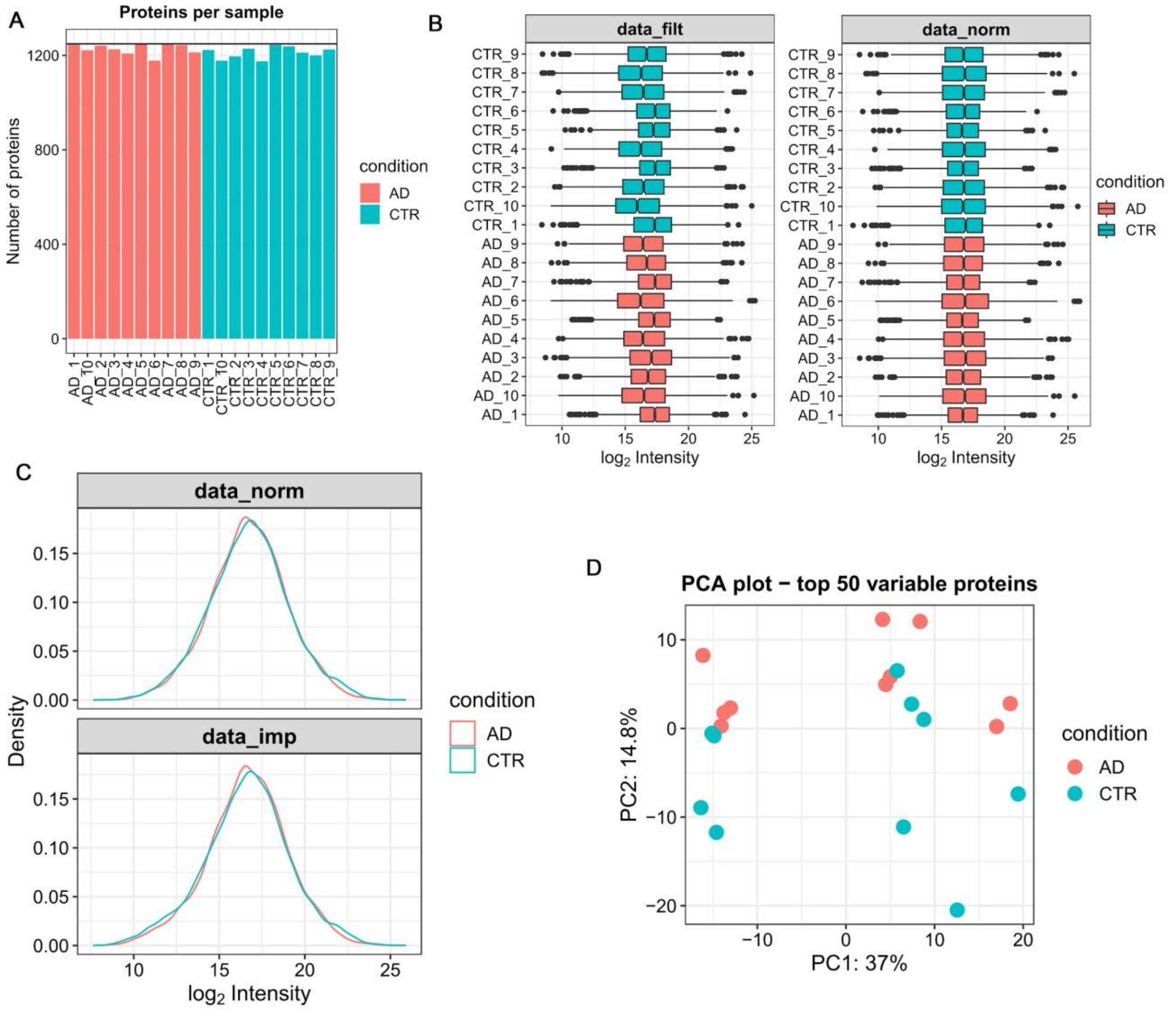


**Supplementary Figure 3**

**Preprocessing Workflow for Purified Proteasome Proteomics from Gray Matter**

(**A**) Bar plot illustrating the total number of proteins detected in each sample. Proteins not identified in all replicates of at least one condition (control or AD) were filtered out prior to downstream processing. (**B**) Box plots showing the distribution of log-transformed peak intensities across samples before and after variance-stabilizing normalization (VSN). This step reduces technical variability in the dataset. (**C**) Density plots comparing log-transformed peak intensities before and after missing data imputation using a left-shifted “MinProb” distribution. Imputation accounts for low-abundance signals absent in the raw data. (**D**) Principal component analysis (PCA) scatter plot using the top 50 most variably expressed proteins. The PCA reveals data structure and checks for potential batch effects among control and AD samples.

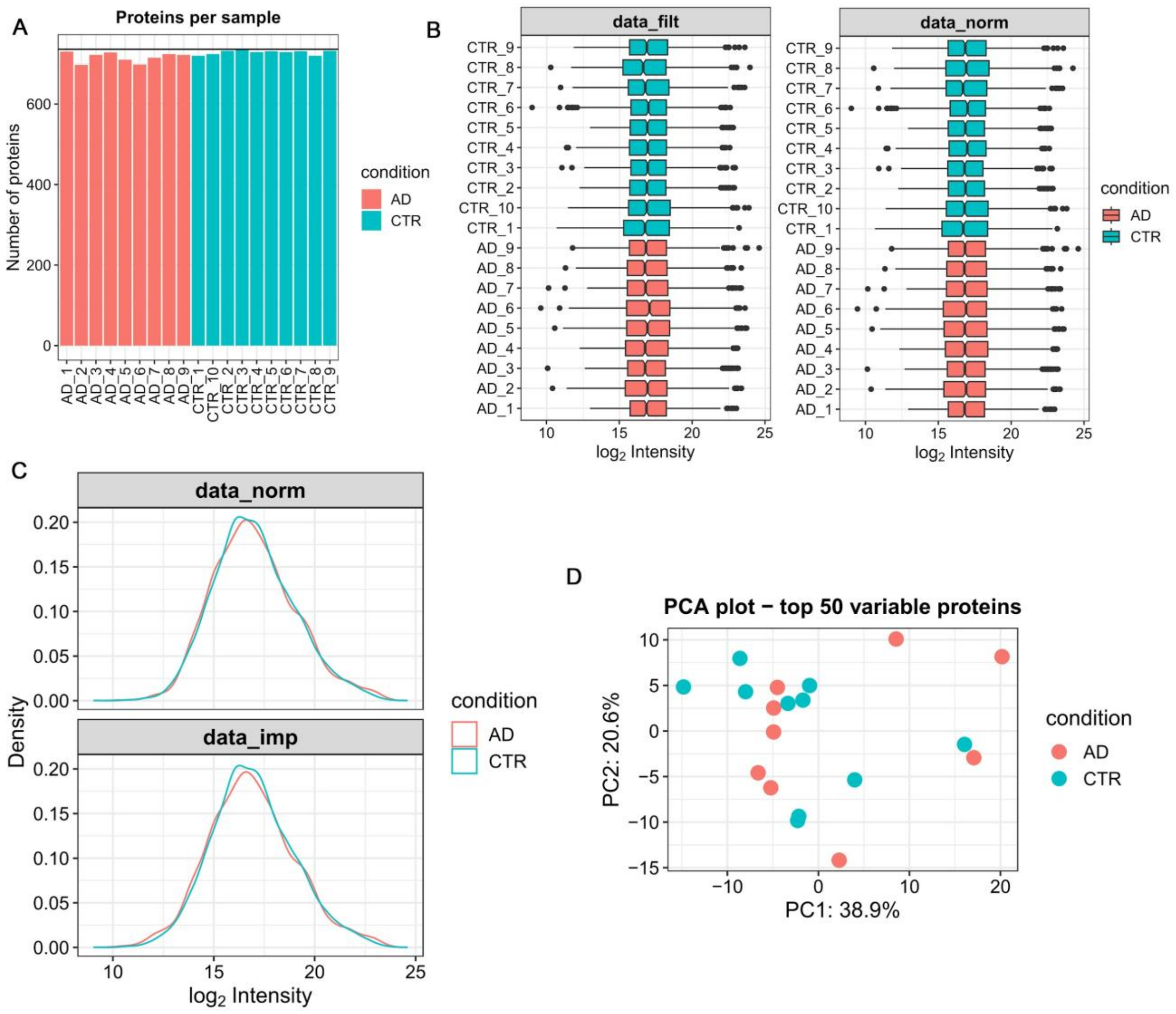


**Supplementary Figure 4**

**Preprocessing Workflow for Purified Proteasome Proteomics from White Matter**

(**A**) Bar plot showing the number of proteins detected per sample. Proteins not identified in all replicates of at least one condition (control or AD) were removed before downstream analyses. (**B**) Box plots of log-transformed peak intensities across samples before and after variance-stabilizing normalization (VSN), demonstrating how VSN helps reduce technical variability. (**C**) Density plots comparing the distribution of log-transformed peak intensities prior to and following missing data imputation with a left-shifted "MinProb" distribution. Imputation compensates for low-abundance signals otherwise not captured in the raw data. (**D**) Principal component analysis (PCA) scatter plot of the top 50 most variably expressed proteins, illustrating the overall data structure and identifying any potential batch effects between control and AD samples.

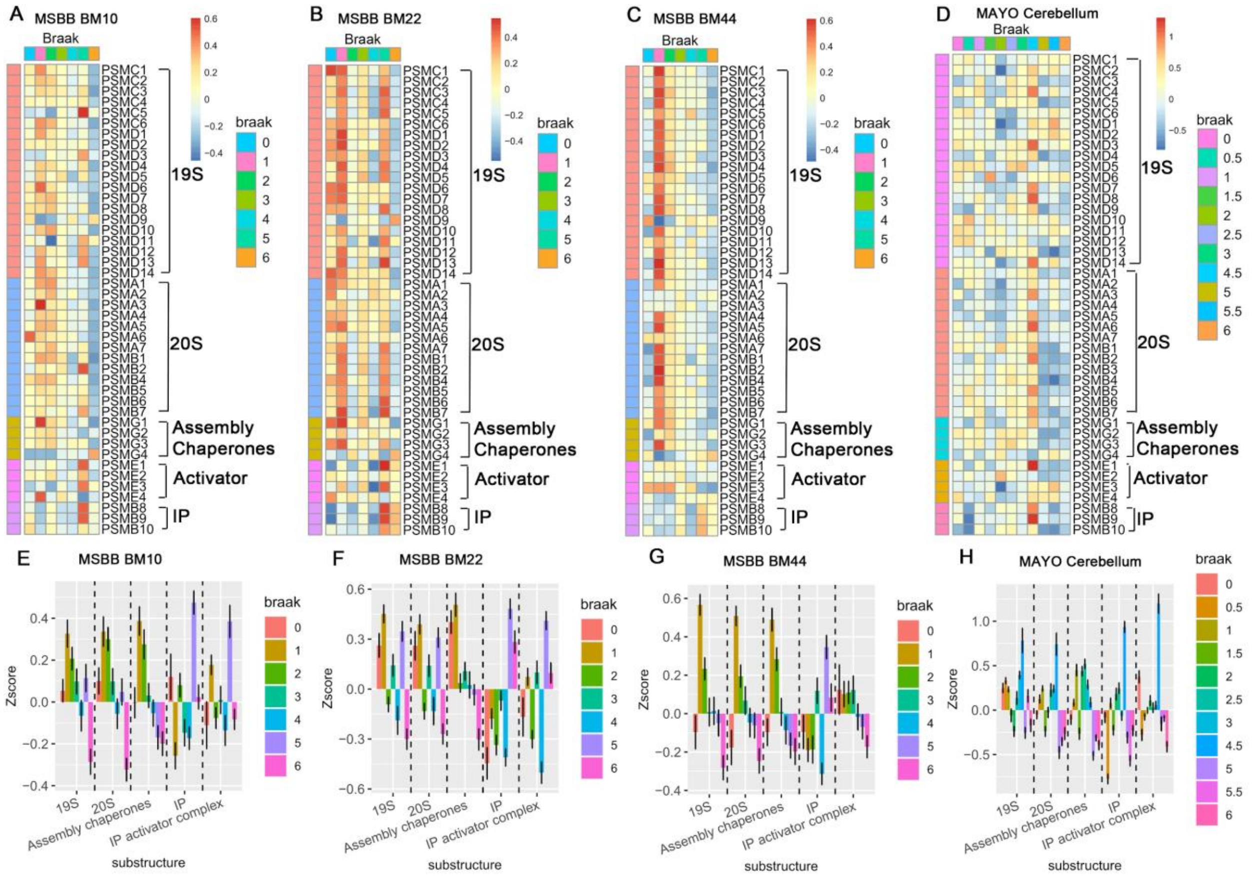


**Supplementary Figure 5**

**Progressive Downregulation of Constitutive Proteasome Subunits and Differential Responses of the Immunoproteasome**

(**A–D**) Heatmaps displaying normalized expression (Z-scores) of constitutive proteasome subunits, proteasome assembly chaperones, and immunoproteasome (IP) components across increasing Braak stages in four different datasets: (**A**) Prefrontal cortex (BM 10); (**B**) Superior temporal gyrus (BM 22); (**C**) Inferior frontal gyrus (BM 44) from the Mount Sinai Brain Bank (MSBB); (**D**) Cerebellum from the Mayo Clinic Study of Aging. Each column represents a sample ordered by Braak stage (0–VI), while each row corresponds to an individual proteasome-related gene. Warmer colors (reds) indicate higher relative expression; cooler colors (blues) indicate lower relative expression. (**E–H**) Box plots of Z-scores for aggregated proteasome complexes and factors—19S, 20S, assembly chaperones, immunoproteasome (IP), and IP activator complexes—grouped by Braak stage in: (**E**) Prefrontal cortex (BM 10); (**F**) Superior temporal gyrus (BM 22); (**G**) Inferior frontal gyrus (BM 44) from MSBB; (**H**) Cerebellum from the Mayo Clinic Study of Aging. These visualizations highlight the pronounced and progressive downregulation of constitutive proteasome genes, contrasted with variable or compensatory changes in immunoproteasome components across different brain regions as AD pathology advances.
